# Targeted Photodegradation of Misfolded Proteins via Self-photosensitizing with Molecularly Produced Light

**DOI:** 10.64898/2026.06.16.732486

**Authors:** Huizhe Wang, Shiju Gu, Jiang Yu, Jinwu Yan, Jing Zhang, Zhiyong Jiang, Caroline Shen, Jun Yang, Chongzhao Ran

## Abstract

Misfolded proteins are tightly associated with various neurodegenerative diseases, and removing these misfolded proteins is one of the actively pursued approaches for seeking therapeutics for these diseases. In this study, we demonstrated that molecularly produced light (molecular light) from ADLumin-5, a self-photosensitizing chemiluminescence compound, could induce photo-oxidation and photodegradation of misfolded proteins, including beta-amyloid, tau, alpha-synucleins, and TDP-43 proteins in vitro. We validated the oxidation and degradation via LC-MS, MADLI-MS, and western blotting. Using beta-amyloid as a showcase, we demonstrated that, upon photo-oxidation and photodegradation, the toxicities of this misfolded protein were significantly reduced. To investigate the therapeutic effects of ADLumin-5 in vivo, we used the 5xFAD mouse model for longitudinal treatment for 4 months. In vivo molecular imaging results indicated that ADLumin-5 could reduce the accumulation of beta-amyloid proteins. Our study presents a novel approach to seek therapeutics for neurodegenerative disease via molecular light-induced degradation of misfolded proteins. In addition, because ADLumin-5 is dual-functional—enabling both photodegradation and in vivo imaging of misfolded protein changes—it can be considered a photo-theranostic agent for neurodegenerative diseases, representing a novel approach to drug discovery for neurodegenerative diseases.

## Introduction

Phototherapy, including UV-, blue, and red-light therapy, photodynamic therapy (PDT), and photo-biomodulation therapy, has been extensively investigated for the treatment of a wide range of diseases, with significant advances achieved in both basic and clinical research ^1–7^. However, the penetration depth of light in biological tissues is inherently limited, restricting the application of phototherapy primarily to superficial targets. Several strategies have been explored to enhance light penetration in vivo, including the use of near-infrared (NIR-I and NIR-II) light, two-photon excitation, upconversion nanoparticles, and interstitial fiber optics ^8–14^. Although these approaches partially mitigate the limitation, they still require external light to reach internal targets, and a substantial proportion of photons is lost during propagation from the external source to the target tissue ^15^.

In previous studies, we and other groups have proposed that molecularly produced light (here we termed as “molecular light”) could be a viable alternative to solve the limitations of light tissue penetration ^15–17^. The most investigated molecular light sources include bioluminescence, chemiluminescence, Cerenkov luminescence, and molecular afterglow ^15,18–24^. In contrast to conventional light sources such as LEDs and lasers, molecular light possesses dual functionality, serving both as a light emitter and as a molecular entity capable of binding to targets. The emitted light can be utilized for phototherapy, while the molecular component enables targeted delivery of the light source to specific biological structures. Upon binding to targets such as proteins, lipids, or glycans, the distance between the light source and the target can approach the zero-to nanometer scale ^15^. At such short distances, emitted photons can be utilized with high efficiency through mechanisms such as energy transfer, direct absorption, or near-zero-distance irradiation. Although the total photon flux generated by molecular light is significantly lower than that produced by conventional light sources, the extremely short interaction distance may compensate for the lower photon output^15,25^.

In 2023, we reported that molecular light from a chemiluminescent compound, ADLumin-4, could be used as a replacement for an external light source for in vivo photodynamic therapy in Alzheimer’s disease (AD) mouse models^25^. In this reported study, we used a bi-molecule combo of photosensitizer (CRANAD-147) and molecular light (ADLumin-4) for the phototherapy study. We demonstrated that ADLumin-4 could photosensitize CRANAD-147 to the initial photo-oxidation of beta-amyloid (Aβ) aggregates. Because this combo therapy involves administering two drug candidates, an obvious drawback is that these two molecules must have similar target binding affinity and PK/PD profiles to form a bio-orthogonal chemiluminescence resonance energy transfer (CRET) pair, necessary to trigger the photo-oxidation reaction with Aβs.

We hypothesized that chemiluminescent compounds with dual functions—both photosensitization and light emission could be used to address this bi-molecule limitation. With these dual functions, the chemiluminescent compound can self-photosensitize, generating ^1^O_2_ upon binding and subsequently initiating photo-oxidizing of Aβs. In addition, this dual function can also serve the purpose of theranostics, which allows for treatment and imaging monitoring simultaneously. Excitingly, indeed, during our continuous studies with the ADLumin-X chemiluminescence series, we discovered that ADLumin-5 was capable of self-photosensitizing as we detected the production of singlet oxygen (^1^O_2_) without external light irradiation. We demonstrated that the ^1^O_2_ generation could initiate the degradation of misfolded proteins, including Aβs, tau proteins, alpha-synucleins, and TDP-43 protein. Interestingly, following our previous studies, in 2024, Umeda et al. reported that a unimolecular chemiluminescence compound alone (dmCLA-X, X = H, Cl, Br) could initiate photo-oxidation of Aβs ^26^. Their strategy was to introduce halogen atoms and -COOH to enhance intersystem conversion (ISC) of the chemi-excited state to the triplet state. However, this strategy is different from our dual-function strategy, as dmCLA-Xs could not be used for theranostics because of the weak chemiluminescence emission of dmCLA-Xs.

Considering PROTAC and Lyso-TAC are highly active areas for drug discovery ^27,28^, our approach of <u>de</u>gradation with <u>mo</u>lecular light (DEMOL) could open an alternative avenue for degrading unwanted proteins (here misfolded proteins in neurodegenerative diseases). Like PROTAC, DEMOL uses small molecules to remove the “bad” unnecessary proteins, instead of inhibiting or agonizing the targets. We believe that phototherapy with the DEMOL platform represents a new, exciting direction and has great potential as an alternative approach to conventional phototherapy.

## Results

### ADLumin-X was capable of self-photosensitizing

To initially evaluate whether molecular light could function as a light source for photosensitization, methylene blue (MB), a well-established photosensitizer ^29^, was used as a model system. Given our previous report showed that ADLumin-5 is highly efficient for chemiluminescence emission ^30^, we first performed tests with ADLumin-5. Indeed, we found that, without external light, the mixture of MB and ADLumin-5 could generate singlet oxygen (^1^O_2_), as detected using 9,10-anthracenediyl-bis(methylene) dimalonic acid (ADPA) ^31^, a commonly used probe for ^1^O_2_ detection (Fig. 1a). After 60 minutes of incubation under dark conditions, approximately a 23.0% decrease in ADPA absorbance at 400 nm was observed, confirming that ADLumin-5 can function as a light source to photosensitize MB. Unexpectedly, a ∼6% decrease in ADPA absorbance was also detected in the control group containing only ADLumin-5. This observation suggested that ADLumin-5 may possess an intrinsic self-photosensitizing capability. In other words, ADLumin-5 appears to exhibit dual functionality, serving as both a light source and a photosensitizer. To determine whether photosensitization is a general property of chemiluminescent compounds, several ADLumin-X analogues (X = −1, −4, −5, −6, −9, −10) were evaluated under LED irradiation, a standard approach for testing photosensitizers. Among these compounds, ADLumin-5 exhibited the highest photosensitizing activity, resulting in an approximately 24% decrease in ADPA absorbance, whereas ADLumin-6 showed the second-highest activity (14% decrease). In contrast, ADLumin-9 did not exhibit detectable photosensitizing activity (Fig. 1b). These results indicate that photosensitizing capability is not a generic property of chemiluminescent compounds but is instead strongly dependent on molecular structure. Although ADLumin-5 is the most effective photosensitizer among the ADLumin-X analogues, it exhibited a lower photosensitizing efficiency, compared to MB (24% vs 75%; SI Fig. 1a). In Fig. 1a, only a modest decrease (∼6%) in the ADPA signal was observed at a relatively low ADLumin-5 concentration and a short incubation time. To determine whether stronger self-photosensitizing effects could occur under extended conditions, the experiment was repeated using 100 μM ADLumin-5 with 4 hours of incubation in the dark. Under these conditions, a pronounced decrease (∼70%) in ADPA absorbance was observed (Fig. 1c), indicating that the self-photosensitizing effect can be both sustained and substantial. Taken together, our results suggested that ADLumin-X, particularly ADLumin-5, was able to self-photosensitize. To the best of our knowledge, this is the first of its kind of self-photosensitizing of small molecules.

**Fig. 1.**
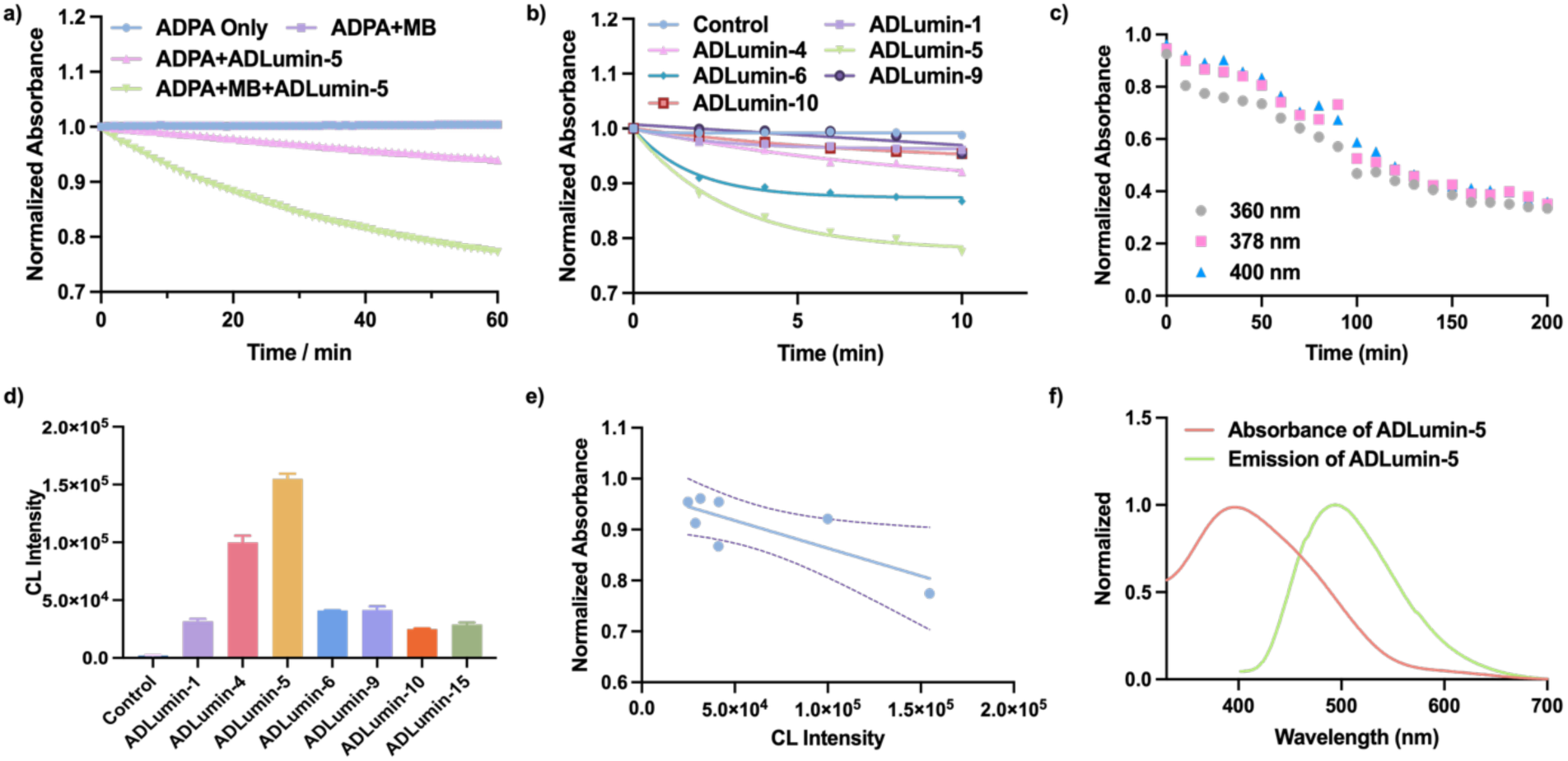
**a**) The detection of singlet oxygen (^1^O_2_) with ADPA (25 μM) during the chemiluminescence generation from ADLumin-5 (25 μM). **b**) ^1^O_2_ generation of ADLumin-Xs (25 μM) incubated with ADPA (25 μM) after LED irradiation; **c**) ^1^O_2_ generation of ADLumin-5 (100 μM) incubated with ADPA (100 μM) in dark condition; **d**) Chemiluminescence intensity of ADLumin-Xs (25 μM); **e**) Correlation fitting of singlet oxygen generation capability and chemiluminescence intensity for ADLumin-Xs; **f**) Spectrum overlap of ADLumin-5 (absorbance and emission).

The ^1^O_2_ generation (self-photosensitizing) capacity of ADLumin-Xs is likely due to the following contributions. First, ADLumin-X is self-illuminating via self-absorption, a common phenomenon for numerous dyes when the absorption/excitation spectrum significantly overlaps with its emission spectrum ^32–34^. In such a scenario, the ^1^O_2_ production efficiency is dependent on the strength of illumination and the overlap of emission and absorption spectra. Indeed, as ADLumin-5 showed the highest ^1^O_2_ capacity, it seems fit in this scenario because it could provide the strongest illumination among the tested ADLumin-Xs (Fig.1d) and there is a linear relationship between chemiluminescence intensity and singlet oxygen yield (Fig. 1e). Furthermore, there is a broad spectral overlap of emission and absorption (Fig.1f). Similarly, spectral overlap also exists between ADLumin-5 and MB (SI Fig. 1b). The second possible contribution is from the chemi-excited state of ADLumin-X, which can be converted into triplet states to initiate the sensitizing of ^3^O_2_ to ^1^O_2_ (Type-II).

The above results indicate that ADLumin-X compounds may function as Type-II photosensitizers, as ^1^O_2_ generation was detected. However, it is also possible that ADLumin-X compounds can operate through a Type-I photosensitization pathway. To investigate this possibility, dihydroethidium (DHE) was employed to monitor superoxide radical generation during the self-photosensitizing reactions of ADLumin-X compounds ^35^. Notably, ADLumin-1 and ADLumin-5 exhibited clear increases in fluorescence intensity, indicating the formation of radical species (SI Fig. 1c). Interestingly, in contrast to the results observed for ^1^O_2_ generation, ADLumin-1 demonstrated a stronger Type-I photosensitizing capability than ADLumin-5. Therefore, the linear relationship between chemiluminescence intensity and superoxide anion yield is also not evident (SI Fig. 1d). These findings also reveal differences compared with a previous report from the Kanai group ^26^. In that study, no detectable ^1^O_2_ generation was observed when furfuryl alcohol (FFA) was used as the detection probe. Instead, hydroperoxyl radicals (HOO•) were proposed as the primary reactive oxygen species responsible for initiating photo-oxidation.

### ADLumin-5 photo-oxidized and degraded model misfolded peptides

In our previous studies, we demonstrated that photo-oxidation could alter the physical and structural properties of Aβs, evident by data from the mass spectrum, SDS-page gel, protein digestion with proteinase K, and seeding activity. Recently, we also found that ADLumin-X could be a generic imaging probe for pan-misfolded proteins, such as aggregates of Aβs, tau proteins, α-synucleins, and TDP-43^36^. The common feature of these misfolded proteins is that all of them are β-sheet aggregates. Based on these results, we reasoned that ADLumin-5 could be applied to a variety of misfolded proteins. In this regard, we first used model peptides that represent β-sheet-enriched misfolded proteins to explore its photo-oxidation capacity. Study results from the Stupp group and our group demonstrated that model peptide PA-K2 (Palmitic-VVVAAAKKK) could form fibrils via the aggregation of β-sheets. Considering our previous studies and other groups’ results, which suggested that the photo-oxidation of Aβ was the oxidation of methionine-35 (M-35), we introduced methionine into the above PA-K2 model peptide by adding it to the middle of Palmitic-VVVMAAAKKK (termed as PAM-K2) to investigate the photo-oxidation (Fig. 2a). To confirm whether PAM-K2 retains the ability to form fibrils, thioflavin T (ThT), a widely used fluorescence probe for detecting β-sheet-rich protein aggregates, was employed to monitor peptide aggregation. As expected, the fluorescence intensity increased by approximately 7.26-fold relative to monomeric PAM-K2 after incubation for three days (Fig. 2b and SI Fig. 2a), indicating that the introduction of methionine did not disrupt the peptide’s aggregation capability. Furthermore, a 167-fold increase in chemiluminescence intensity was observed upon mixing ADLumin-5 with PAM-K2 fibrils, suggesting that ADLumin-5 can effectively bind to the fibrillar aggregates (Fig. 2c and SI Fig. 2b).

**Fig.2.**
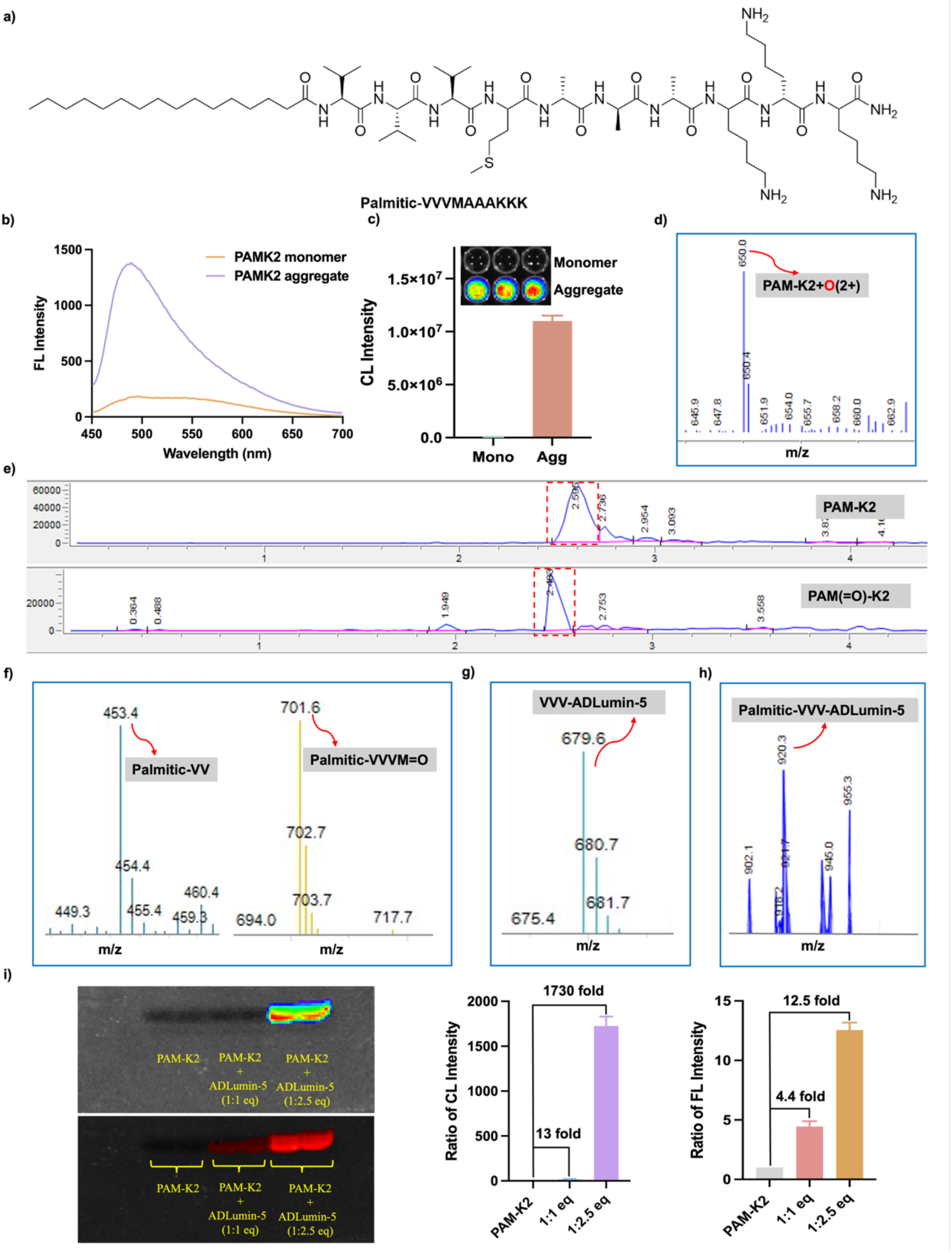
**a)** Structure of Palmitic-VVV**M**AAAKKK (PAM-K2); **b)** fluorescence spectrum of PAMK2 monomer or aggregate (5 μM) after added Thioflavin T (0.5 μM) in PBS; **c)** Chemiluminescence imaging of PAMK2 monomer or aggregate (50 μM) with ADLumin-5 (50 μM); **d)** LC-MS analysis of PAM-K2 with ADLumin-5; **e)** HPLC analysis of PAM-K2 before and after oxidize; **f)** Cleavaged fragment of PAM-K2 after oxidized; **g-h)** PAM-K2 fragment conjugated with ADLumin-5; **i)** SDS-PAGE of PAM-K2 aggregate (20 μM) with or without ADLumin-5.

To investigate whether ADLumin-X can induce photo-oxidation of PAM-K2, we used PAM-K2 peptide itself to run LC-MS as the control group, from which we observed m/z = 642.2 at 2.63 minutes, which is corresponding to the MS of PAM-K2 peptide with 2 charges, and m/z=1282.6 for PAM-K2 peptide (SI Fig.2c). Then we incubated ADLumin-5 with the PAM-K2 aggregates for 24 hours and used LC-MS to identify the products. Indeed, we found several MS peaks that were related to the oxidation. First, we identify m/z = 650.2 (2+) as the oxidized form of PAM-K2 (+16) (Fig.2d), suggesting that ADLumin-5 indeed can photo-oxidize this model peptide. As we expected, the oxidized PAM(=O)-K2 showed larger polarity, evident by its shorter retention time (2.48 minutes vs 2.63 minutes) (Fig.2e). The ratio of PAM-K2/PAM(=O) was about 3:1. In addition, a peak at m/z = 702 was detected (Fig. 2f), which was assigned to Palmitic-VVVM(=O). This observation indicates that methionine oxidation occurred simultaneously with peptide fragmentation, suggesting that ADLumin-5 can mediate both photo-oxidation and peptide degradation. Furthermore, a fragment with m/z = 453.3 corresponding to Palmitic-VV was also observed (Fig. 2f), providing additional evidence that ADLumin-5 can induce degradation of the PAM-K2 peptide. Unexpectedly, a strong peak at m/z = 679.3 with a retention time of 1.84 min was detected (Fig. 2g), indicating the formation of a highly polar product. Based on detailed mass analysis, this peak was assigned to VVV-ADLumin-5, representing a conjugation product between ADLumin-5 and the VVV peptide fragment. In addition, a moderate peak at m/z = 920.3 with a retention time of 3.03 min was detected (Fig. 2h) and assigned to Palmitic-VVV–ADLumin-5, further supporting the formation of ADLumin-5–peptide conjugates.

To further validate the conjugation of ADLumin-5 to peptide fragments, SDS-PAGE analysis was performed to determine whether fluorescence and chemiluminescence signals could be detected, as ADLumin-5 possesses both fluorescent and chemiluminescent properties. Consistent with this expectation, a distinct band exhibiting strong fluorescence and chemiluminescence signals was observed (Fig. 2i), supporting the formation of ADLumin-5–peptide conjugates. Collectively, these results indicate that ADLumin-5 can induce photo-oxidation, photo-degradation, and photo-conjugation of PAM-K2 aggregates. The oxidation events are consistent with the modification of the methionine residue within the peptide sequence. We also performed tests with all the ADLumin-Xs with PAM-K2 and calculated their ratios of oxidized/non-oxidized. Among all ADLumin-Xs, ADLumin-5 showed the best results in photo-oxidation (SI Table 1). Based on this result, in the following studies, only ADLumin-5 was used for a comprehensive investigation.

### ADLumin-5 photo-oxidized and degraded Aβs

Our previous study with a combo of photosensitizer and ADLumin-4 and Kanai group’s recent unimolecular results showed that Aβs could be oxidized with molecular light, and identified that M-35 is the amino acid to be photo-oxidized ^25,26^. Our above results with the model peptide again showed that M-35 could be oxidized by molecular light. To investigate whether self-photosensitizing of ADLumin-5 can oxidize Aβs, we performed a series of experiments. First, we used high-resolution LC-MS to identify the possible products after incubation of ADLumin-5 and Aβs for 24 hours. As we expected, m/z = 4345.1596, which corresponds to oxidized Aβ_40_ with an addition of one oxygen, could be clearly observed, and this peak intensity was even higher than the Aβ_40_ peptide itself (m/z = 4329.1632) (Fig. 3a). This result unambiguously suggested that ADLumin-5 alone could photo-oxidize Aβs. Excitingly, we also observed m/z = 3786.8792, which corresponds to the molecular weight of Aβ_1-34_, suggesting ADLumin-5 could degrade Aβ40 (Fig. 3a). Notably, we also observed m/z = 4197.8929, which is likely corresponding to Aβ_1-34_+ADLumin-5+O (Fig. 3a). This result suggested that ADLumin-5 could form covalent conjugation with Aβ fragments during the photo-reaction. To further validate the observed photo-oxidation and degradation, we further used LC-MS to analyze the Aβ fragments after trypsinization, which is specific to lysine (K) and arginine (R) cleavage. The K/R specific cleavages could be easily found from the control, including Aβ_1-5_, Aβ_6-16_, Aβ_17-28_, and Aβ_29-40_ (SI Fig. 3a), suggesting the trypsinization is effective. As expected, oxidized fragments, such as Aβ_29-40_+O, could be identified. Excitingly, we observed several specific fragments that contain K but not R within the sequence, suggesting that the fragments are not from trypsinization. For example, we found m/z = 801.3 peak, which corresponds to Aβ_6-34_+2O (4+charges). We also observed m/z = 1237.7 peak, which could likely be assigned to a trimer of Aβ_17-34_ with one oxygen addition (3+charges) (Fig. 3b). Again, the LC-MS data further support that molecular light from ADLumin-5 can degrade Aβs at the L-34 position, which is close to M-35. We also performed similar experiments with Aβ_42_ aggregates and found similar results (SI Fig. 3b). The above MS results suggested that Aβ peptides could be oxidized, degraded, and conjugated during the reactions with ADLumin-5. To further provide solid evidence, we used SDS-PAGE to investigate the changes in Aβ peptides. First, we used a fluorophore-tagged Aβ peptide (FAM-Aβ_42_) to perform the electrophoresis. As expected, a degraded band could be clearly observed in the presence of ADLumin-5 (Fig. 3c). We also observed a similar degraded band with FAM-Aβ_40_ aggregate (SI Fig. 3c). This is consistent with our evidence from MS studies. Encouraged with this result, we performed western blots with native Aβs (without fluorophore tag) with Aβ antibody 6E10. As expected, a band with molecular weight < 4 KD could be clearly observed for both Aβ_40_ and Aβ_42_ aggregates (Fig. 3d), again suggesting ADLumin-5 can induce degradation of Aβs via self-photosensitization.

**Figure 3.**
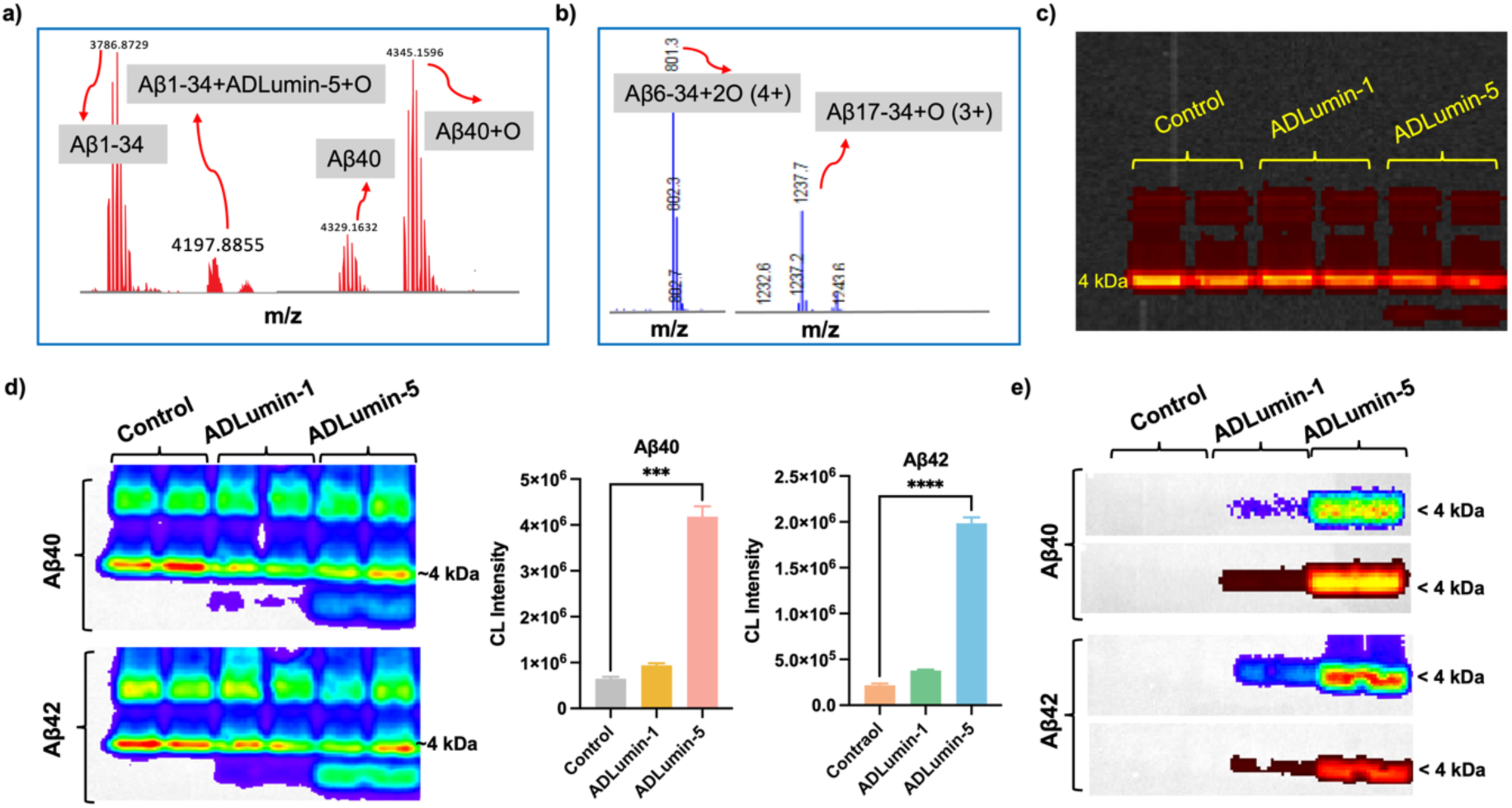
**a**) High resolution LC-MS analysis of Aβ40 incubated with ADLumin-5; **b**) LC-MS analysis of Aβ40 incubated with ADLumin-5 after trypsin cleavage; **c**) SDS-PAGE of FAM-Aβ42 aggregate incubated with ADLumin-Xs; **d**) Western blot assay (after antibody incubation) and quantification of Aβ aggregate (20 μM) incubated with different ADLumin-Xs (20 μM); **e**) Western blot assay (before antibody incubation) of Aβ aggregate (20 μM) incubated with different ADLumin-Xs (20 μM), image were obtained in both chemiluminescence and fluorescence channel. Data are mean ± SD, n = 3. ***p < 0.001, ****p < 0.0001

Notably, as with the observations obtained with the PAM-K2 model peptide, a band exhibiting both chemiluminescence and fluorescence signals was detected before antibody incubation. This band migrated faster than the monomeric Aβ band (Fig. 3e), suggesting that it corresponds to a degradation product conjugated with ADLumin-5. This interpretation is consistent with the MS signal at m/z = 4197.8929, which was assigned to Aβ_1-34_ conjugated with ADLumin-5 and one oxygen atom (Aβ_1-34_+ADLumin-5+O). Further analysis revealed that the formation of this conjugation–degradation band, which could be detected through both chemiluminescence and fluorescence signals from ADLumin-5, was dependent on both ADLumin-5 concentration and incubation time (SI Fig. 3d). It is worth noting that the monomeric band, as well as other bands, may contain a high proportion of oxidized Aβ species, as indicated by our MS analyses. Taken together, data from MS, SDS-page, and western blots indicated that ADLumin-5 could induce photo-oxidation, degradation, and conjugation of Aβs without external light irradiation. The results from Aβs are consistent with studies with the PAM-K2 model peptide, suggesting that such photo-oxidation and photo-degradation could be a generic phenomenon for misfolded proteins.

### ADLumin-5 selectively oxidized the Met35 residue in Aβ peptides

To identify the specific peptide fragments and oxidation sites in Aβ, the degradation band of Aβ_40_ aggregate treated with ADLumin-5 after SDS-PAGE was subjected to LC–MS/MS analysis. Interpretation of the tandem mass spectra revealed that the primary oxidized fragment corresponded to Aβ_29-40_ (sequence: GAIIGLMVGGVV). The doubly charged precursor ion ([Aβ_29-40_ + 2H]²⁺) was detected at m/z = 551.25, corresponding to a molecular weight of 1101.63 Da (SI Fig. 5). The MS/MS spectrum exhibited a series of consecutive b and y fragment ions that closely matched the theoretical fragmentation pattern of Aβ_29-40_. Notably, the b_7_^+^ ion peak shifted from m/z 656.88 (unoxidized) to m/z 672.38, corresponding to a mass increase of approximately 16 Da, which indicates oxidation of the seventh residue in this fragment (methionine, Met35) to methionine sulfoxide (Met(O)). Consistently, the y_6_^+^ ion peak shifted from m/z 561.72 to m/z 577.30, further confirming oxidation at the Met35 site (SI Tab. 2 and Tab. 3). These MS/MS results are consistent with the peptide matching analysis and confirm that ADLumin-5–mediated oxidation of Aβ predominantly occurs at the Met35 residue. Methionine residues are highly susceptible to oxidation by ^1^O_2_ or other ROS, forming methionine sulfoxide. Accordingly, ADLumin-5 is proposed to generate ROS through self-photosensitization, leading to the selective oxidation of Met35 in Aβ. Oxidation of this residue is expected to alter the hydrophobicity and conformational stability of the peptide fragment, which may disrupt the β-sheet structure and contribute to the structural destabilization and subsequent degradation of Aβ aggregates.

### ADLumin-5 selectively oxidized β-sheet–rich proteins

To investigate whether the photo-oxidation is β-sheet–rich specific, we compared the fluorescence signals generated after incubating ADLumin-5 with Aβ peptides and with non-amyloid (without β-sheet) model peptides containing oxidation-sensitive amino acid residues. As shown in SI Fig. 6a, ADLumin-5 exhibited the strongest fluorescence signal when incubated with Aβ_42_ aggregates, indicating the highest binding affinity, followed by Aβ40 aggregates. In contrast, incubation with Aβ_40_ and Aβ_42_ monomers, as well as with non-β-sheet proteins (such as Angiotensin IV, NKA, and RNase A^26^), produced relatively weak and comparable fluorescence signals. These results suggest that ADLumin-5 exhibits a clear structural preference for β-sheet–rich conformations, with particularly enhanced recognition of aggregated Aβ species. Consistent with this observation, the oxidation yield of ADLumin-5 toward Aβ_40_ aggregates was 119-fold higher than that toward Angiotensin IV, while the oxidation yield toward Aβ_42_ aggregates was 85-fold higher than that toward Angiotensin IV (SI Fig. 6b).

### ADLumin-5 altered properties of Aβs

After the photo-oxidation, we speculated that Aβ’s properties, such as aggregating tendency and resistance to proteinase K, could be altered. In this regard, we first performed a binding recovery study, in which we incubated Aβ aggregates with ADLumin-5 for 24 hours and then added ADLumin-5 again. We expected the freshly added ADLumin-5 would not result in significant fluorescence intensity increases, since the aggregates were destroyed by the photo-oxidation and degradation. Indeed, no significant increase in fluorescence was observed after the addition of the fresh ADLumin-5 (Fig. 4a). We didn’t use thioflavin T (ThT) to monitor the alterations, because the fluorescence spectra of ADLumin-5 and ThT have a strong overlap. Moreover, we investigated whether the photo-oxidation could accelerate digestion of Aβs with proteinase K. To this end, we compared the time-course of fluorescence change of ADLumin-5 in the presence of proteinase K. We found that the ADLumin-5 treated group (24 hours incubation) had much fast decay of fluorescence, compared to the control group (without ADLumin-5 pre-treatment) (Fig. 4b). Moreover, we performed seeding experiments with Aβ aggregates that were treated with ADLumin-5 for 24 hours, and the fluorescence of ADLumin-5 was used to monitor the aggregation process. Compared to the control groups (no seeds that contained PBS buffer only and Aβ seeds that were not treated with ADLumin-5), we found that the seeds from the ADLumin-5-treated group showed much slower aggregations (Fig. 4c).

**Fig 4.**
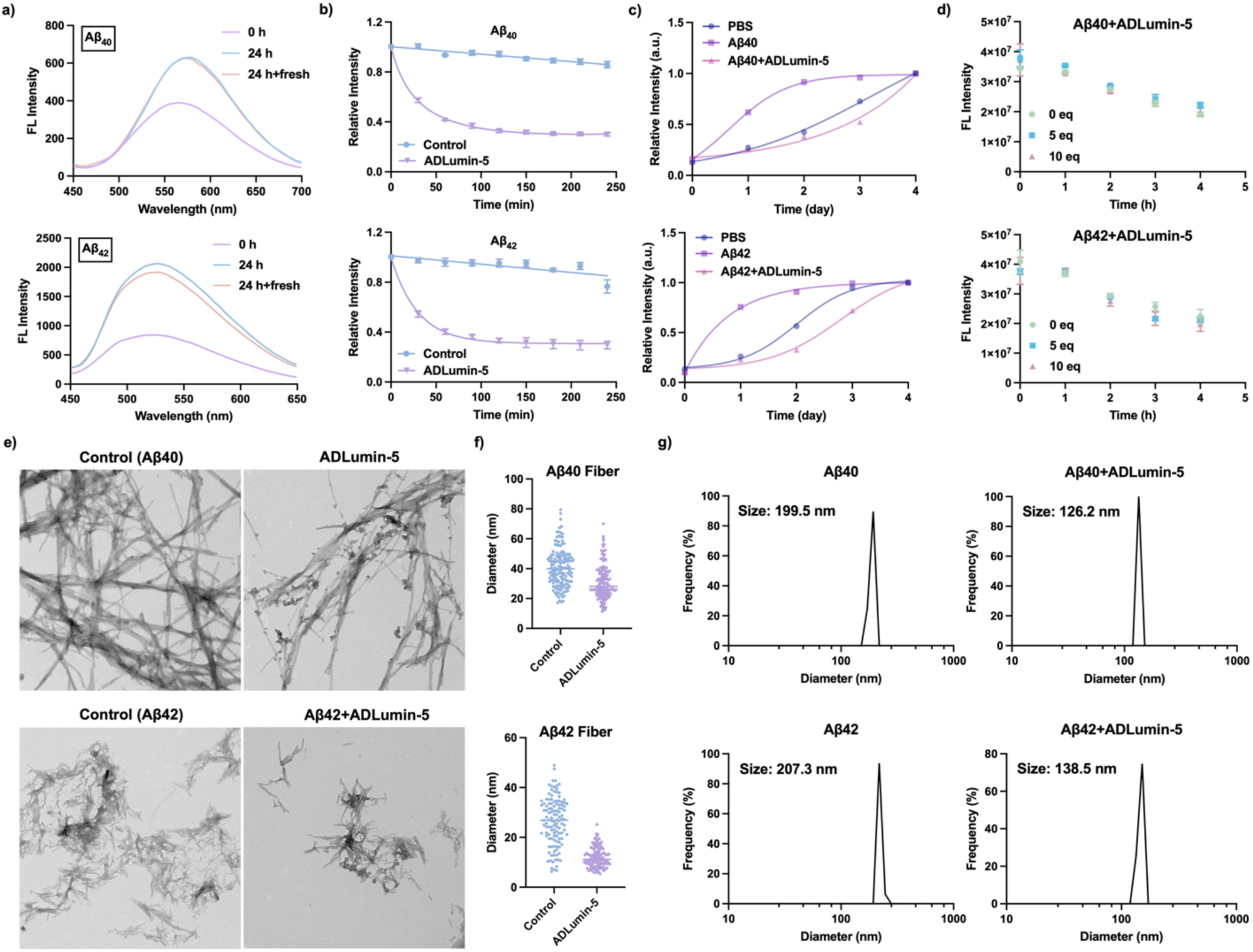
**a**) Recovery study of Aβ (2μM) with different treatments; **b**) Time-course of the degradation of Aβs (2 µM) with proteinase K (0.1 mg/mL); **c**) Relative fluorescence intensity and time-course of Aβ aggregation in the seeding experiments; **d**) Fluorescence intensity of Aβ aggregate in the presence of ADLumin-Xs with different concentrations of catechol at different time points; **e**) TEM imaging of Aβ morphology changes with or without ADLumin-5 incubation; **f**) Diameter quantification of Aβ fibers with or without ADLumin-5; **g**) Dynamic laser scattering size of Aβ with or without ADLumin-5

Given the potential concern regarding the neurotoxicity of ADLumin-5 during its therapeutic action, we investigated whether the photo-oxidation process is largely confined within Aβ aggregates, thereby minimizing off-target reactions. To evaluate this possibility, catechol was employed as a cocatalyst because it has been reported to enhance photon-induced free-radical reactions while exhibiting negligible binding affinity toward Aβ aggregates ^37^. Under this design, if reactive oxygen species were to diffuse away from the Aβ aggregates, they would be expected to react with catechol and generate downstream oxidative products. As anticipated, the fluorescence intensity of ADLumin-5 decreased after its addition to the mixture. Importantly, no acceleration of fluorescence decay was observed in the presence of catechol (Fig. 4d), suggesting that catechol did not interact with reactive oxygen species to promote additional radical formation. These results indicate that the photo-oxidation process mediated by ADLumin-5 is likely confined to the vicinity of Aβ aggregates, thereby limiting oxidative reactions outside the aggregates.

Transmission electron microscopy (TEM) was also used to examine the morphology of Aβ fibrils with and without ADLumin-5 treatment. The fibrils formed after incubation with ADLumin-5 displayed a substantially smaller diameter compared with untreated fibrils (Aβ_40_, 31 nm vs 41 nm; Aβ_42_, 12 nm vs 26 nm) (Fig. 4e and Fig. 4f), further indicating structural alterations induced by ADLumin-5. Further measurements of the hydrodynamic diameters of the aggregates by dynamic light scattering (DLS) revealed that, following photo-oxidation, the average diameter of Aβ_40_ aggregates decreased from 199.5 nm to 126.2 nm, while that of Aβ_42_ aggregates decreased from 207.3 nm to 138.5 nm (Fig. 4g). These results indicate that the photo-oxidative activity of ADLumin-5 disrupts the polymeric aggregated state of Aβ, leading to the formation of smaller oligomeric species. The zeta potentials of untreated Aβ_40_ and Aβ_42_ oligomers were −51.6 mV and −45.4 mV, respectively, whereas incubation with ADLumin-5 reduced the absolute values to −37.0 mV and −28.8 mV, respectively (SI Fig. S4). This change suggests that photo-oxidation by ADLumin-5 alters the oxidation states of surface-exposed amino acid residues in Aβ, thereby modifying the surface charge distribution and intermolecular interactions. Such alterations likely weaken the electrostatic stability of the aggregates, ultimately promoting their depolymerization and degradation.

### ADLumin-5 photo-oxidized and degraded other misfolded proteins

Previous studies have shown that ADLumin-1 serves as a generic imaging probe for a range of misfolded proteins ^36^, including Aβ, tau, α-synuclein, and TDP-43 ^38–40^. The model peptide experiments described above further indicated that ADLumin-5 is capable of inducing photo-oxidation and degradation of β-sheet–rich structures. Based on these findings, it was hypothesized that ADLumin-5 could also act on other misfolded proteins. To test this possibility, ADLumin-5 was incubated with preformed fibrils of α-synuclein for 24 h. A degraded band exhibiting both chemiluminescence and fluorescence signals was clearly observed on SDS-PAGE gels. Compared with the chemiluminescence signal, the fluorescence signal was more pronounced (Fig. 5a). The molecular weight of the fluorescence band was approximately 4.3 kDa, suggesting cleavage near the α-synuclein 103–140 region (around K_102_). However, this degraded chemiluminescent band was not detectable after Western blotting using a synuclein antibody and luminol-based chemiluminescence imaging, likely due to the weak signal from the degraded fragment and the inability of the antibody to recognize the cleaved sequence, or ADLumin-5 underwent a ring-opening reaction after covalently binding to the degradation band. LC–MS analysis of trypsin-digested α-synuclein fibrils treated with ADLumin-5 for 24 h identified several oxidized fragments, including α-syn_1–6_ +O (m/z = 393.1, 2+) and α-syn_103–140_ +O (m/z = 1443.6, 3+) (Fig. 5b). Both fragments contain methionine residues that are susceptible to oxidation. In addition, a peak at m/z = 1007.2 corresponding to α-syn_1–39_ +O (4+) was observed (Fig. 5c). Because no lysine residue is present near position 39 for trypsin cleavage, this fragment likely corresponds to the degradation band with a molecular weight of approximately 4 kDa observed in the gel analysis.

**Figure 5.**
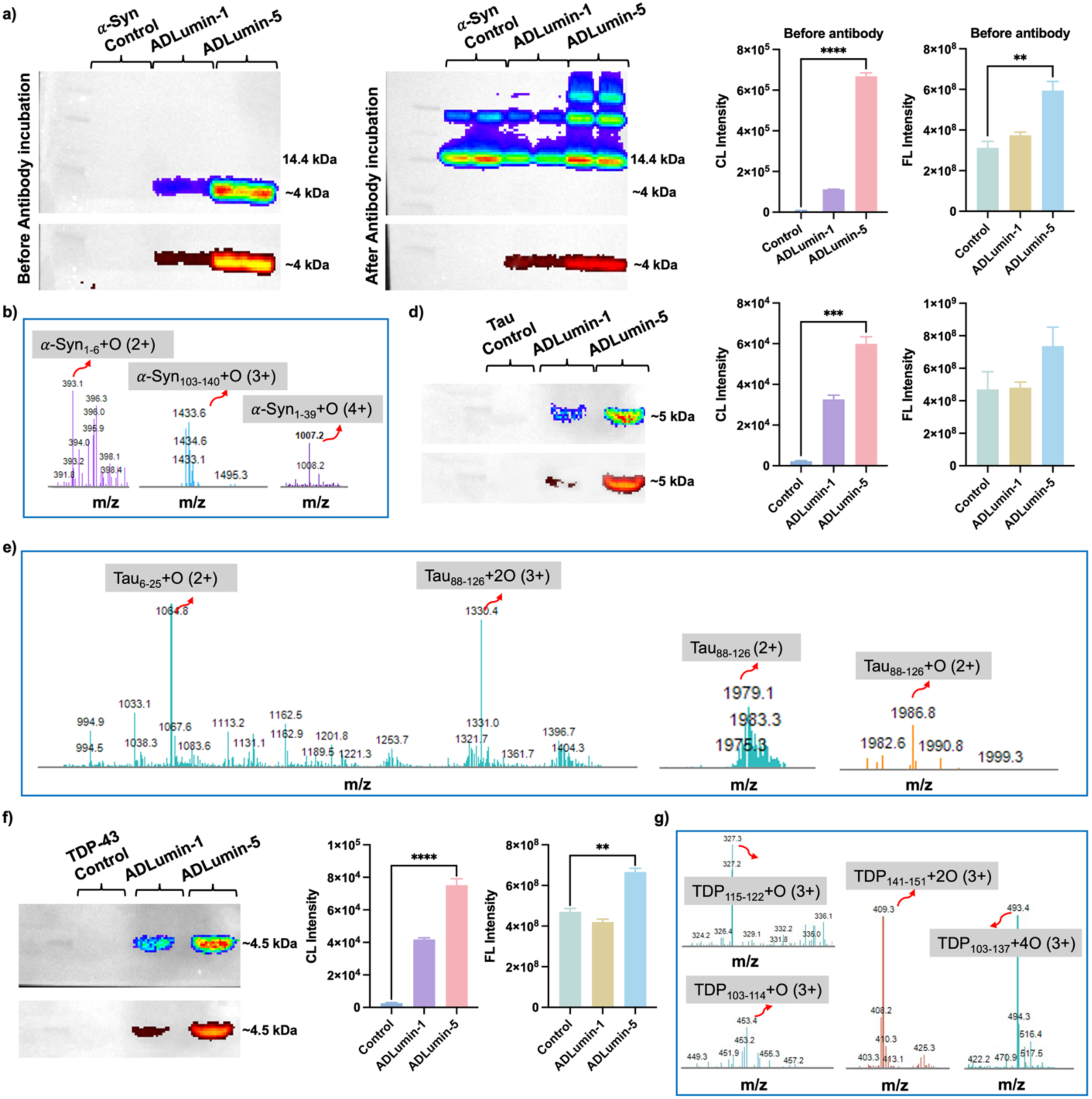
**a**) Western blot of Alpha-Synuclein (*α*-syn) treated with ADLumin-Xs and degradation band quantification before antibody incubation. Data are mean ± SD, n = 3. **p < 0.01, ****p < 0.0001; **b-c**) LC-MS analysis of *α*-Synuclein incubated with ADLumin-5; **d**) Western blot of Tau 441 (2N4R) treated with ADLumin-Xs (before antibody incubation). Data are mean ± SD, n = 3. ***p < 0.001; **e**) LC-MS analysis of Tau-441 (2N4R) incubated with ADLumin-5; **f**) Western blot of TDP-43 treated with ADLumin-Xs (before antibody incubation. Data are mean ± SD, n = 3. **p < 0.01, ****p < 0.0001; **g**) LC-MS analysis of TDP-43incubated with ADLumin-5.

Next, incubation experiments were conducted using preformed fibrils (PFFs) of Tau-441 (2N4R) in the presence of ADLumin-5. Like the results obtained with α-synuclein, a degradation/conjugation band with both chemiluminescence and fluorescence signals was observed, with a molecular weight of approximately 5 kDa (Fig. 5d). LC–MS analysis identified several oxidized fragments, including Tau_6–25_ +2O (m/z = 1064.8, 2+), Tau_88–126_ +2O (m/z = 1330.4, 3+), and Tau_88–126_ +O (m/z = 1986.8, 2+). Based on the Tau sequence, the degradation site is likely located near H_88_ and H_89_, as histidine residues are also susceptible to oxidation ^41,42^ (Fig. 5e).

TDP-43 protein is another well-studied protein that is prone to aggregation and tightly associated with amyotrophic lateral sclerosis (ALS) and frontotemporal lobar degeneration (FLTD) ^40^. With TDP-43 aggregates, we performed SDS-PAGE gel and western blotting after 24-hour incubation with ADLumin-5. Similar to alpha-synucleins and Tau-441 (2N4R), we observed a degraded/conjugated band with a molecular weight of around 4.5 KD (Fig. 5f), and this cleavage is likely related to C39. We also performed LC-MS analysis with TDP-43 aggregate after incubation with ADLumin-5 for 24 h and trypsinization for 18 h. We observed that there were several oxidized fragments, including TDP_103-114_+O with three charges (m/z=453.4), TDP_103-137_+4O with three charges (m/z=493.4), TDP_115-122_+O with three charges (m/z=327.3), and TDP_141-151_+2O with three charges (m/z=409.3). Interestingly, all the degraded bands could be due to the photo-cleavage around N or C terminals. These alterations could be beneficial for reducing the toxicity of these misfolded proteins. Taken together, consistent with the results from the PAM-K2 model peptide and Aβs, molecular light from ADLumin-5 was capable of inducing photo-oxidation, degradation, and conjugation of a variety of misfolded proteins.

### ADLumin-5 attenuated the cytotoxicity induced by Aβs in cell and organoid studies

It has been widely reported that oxidized Aβ species exhibit reduced neurotoxicity compared with their non-oxidized counterparts^25,26,43–46^. The Aβ_25-35_ fragment has been extensively used to induce cytotoxicity in both in vitro and in vivo studies ^47,48^. Notably, oxidation occurs at Met35, which resides within the hydrophobic region of Aβ and plays a critical role in aggregation. It is therefore reasonable to speculate that oxidation at this site attenuates neurotoxicity. Consistent with this notion, our results indicate that degradation also occurs near Met35, potentially disrupting the hydrophobic segment and thereby reducing toxicity. Conceivably, such degradation could also reduce the aggregation and attenuate Aβ’s neurotoxicity. To investigate whether ADLumin-5 can attenuate the cytotoxicity of Aβs, we incubated neuronal cells SH5-SY5 with Aβs (as the control) and oxidized Aβs that were treated with ADLumin-5 before adding them to the cell culture medium. As we expected, compared to the control (the Aβ group), the ADLumin-5 group showed higher ATP concentrations, suggesting that the oxidized Aβs have lower cell toxicity (SI Fig. 7a). To further support this observation, we also performed MTT tests with the same groups. Consistent with the ATP tests, the oxidized Aβ group showed much weaker cell toxicity (Fig. 6a). To further evaluate toxicity under conditions that more closely resemble in vivo environments, neuronal spheroids were cultured and exposed to the same treatment conditions. As expected, spheroids treated with oxidized Aβ displayed reduced toxicity compared with those treated with untreated Aβ aggregates (SI Fig. 7b). Notably, ADLumin-5 itself showed no apparent toxicity with SH5-SY5 cells (SI Fig. 7c).

**Figure 6.**
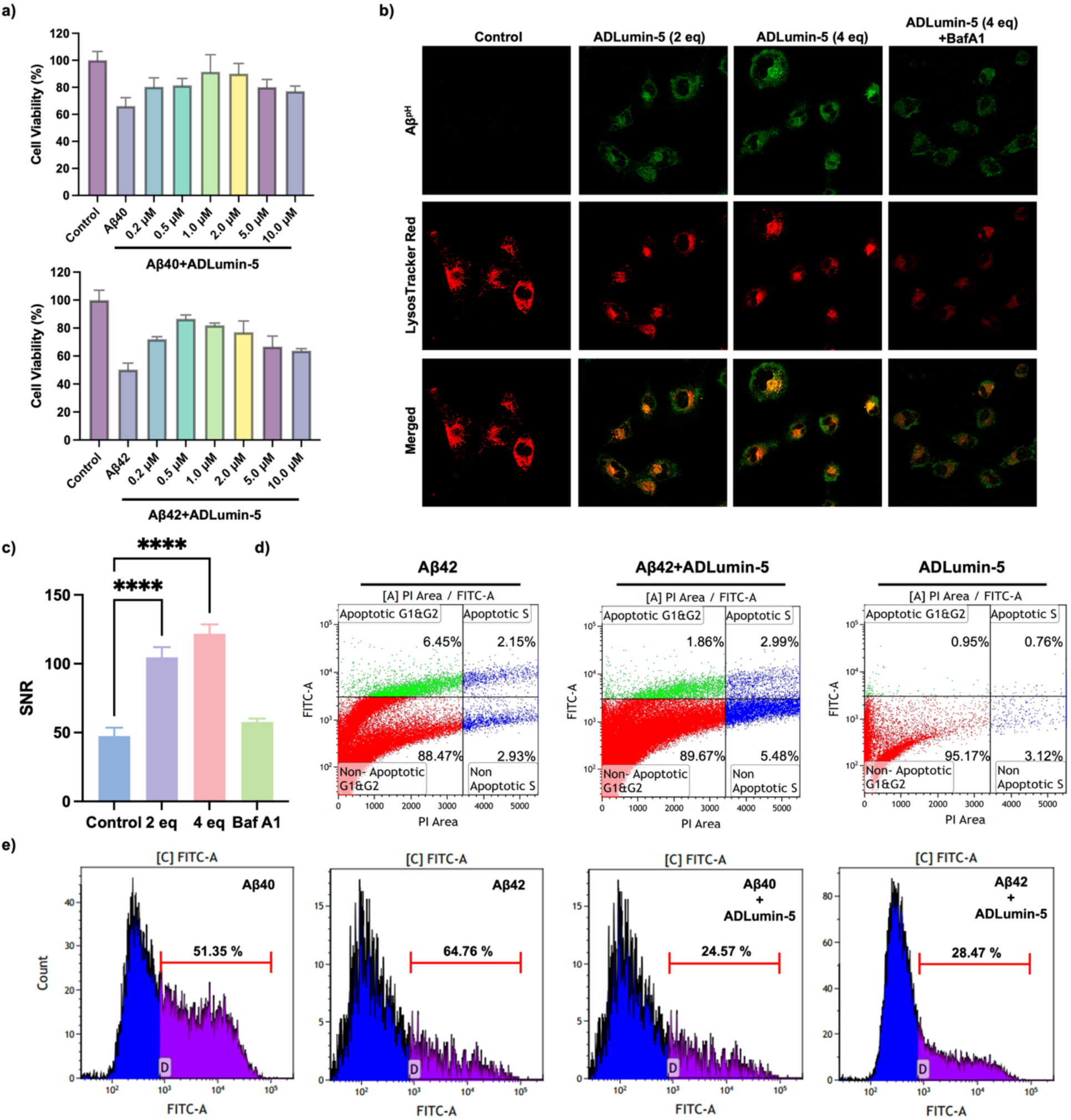
**a**) MTT assay of Aβ (20 μM) treated with different concentrations of ADLumin-5; **b**) The effect of ADLumin-5 on lysosomal uptake of amyloid beta; **c**) SNR study of the green channel. Data are mean ± SD, n = 3. ****p < 0.0001; **d-e**) 3D spheroids and organoids apoptosis study of Aβ with or without ADLumin-5.

It is well-documented that autophagy-regulated uptake and lysosomal degradation of Aβs are essential for reducing neurotoxicity ^49,50^. In this regard, we used the Aβ^PH^ probe to monitor the uptake and trafficking of Aβs after ADLumin-5 treatment ^49^. As expected, higher uptake was observed in the ADLumin-5 groups compared to the control group, and the ADLumin-5-treated Aβs were co-localized with lysoTracker (Fig. 6b,c). To further confirm the autophagy effect, we used Bafilomycin A1 (BafA1), a widely used inhibitor of autophagy and lysosomal acidification ^51^, to perform the experiments. Indeed, BafA1 could significantly inhibit the Aβ uptake, suggesting that, indeed, the oxidized Aβs could be degraded via autophagy (Fig. 6b,c). To further quantify lysosomal colocalization, Pearson’s correlation coefficients between Aβ^pH^ and lysosomal signals were calculated. In the absence of ADLumin-5, the Pearson coefficient was only 0.03. However, after the addition of 10 μM ADLumin-5, the coefficient increased 12-fold, indicating enhanced lysosomal localization of Aβ. Higher concentrations of ADLumin-5 further increased the colocalization coefficient, whereas treatment with BafA1 significantly reduced the value, confirming the critical role of lysosomal activity in the observed fluorescence enhancement (SI Tab. 4).

Moreover, to test the toxicity under conditions more relevant to the in vivo environment, we cultured organoids of neuron cells and incubated them with Aβs or ADLumin-5/ Aβs. As expected, the oxidized Aβ group showed lower toxicity compared to the non-oxidized control group, as evident by less apoptosis from the G1/G2 phase (Fig. 6d). Again, ADLumin-5 itself showed no apparent apoptosis for the phases, indicating ADLumin-5 is not toxic for the organoids. We further confirmed the reduced toxicity of the ADLumin-5 group with the spheroid culture of neuronal cells, as evident by the reduced apoptosis rate from 51.35% (treated with Aβ_40_) and 64.76% (treated with Aβ_42_) to 24.57% and 28.47%, respectively (Fig. 6e). Taken together, these results demonstrate that ADLumin-5-mediated photo-oxidation markedly attenuates Aβ-induced neurotoxicity. This protective effect is reflected not only in improved cell viability but also in reduced apoptosis across multiple cellular and organoid models.

### ADLumin-5 induced Aβ degradation in brain homogenates, brain organoids, and brain tissues

To investigate whether ADLumin-5 can induce degradation in biologically and pathologically relevant environments, we first used wild-type (WT) mouse brain homogenates to mimic such environments. We treated brain homogenates that were added to Aβ_40_ aggregates with ADLumin-5 for 24 hours, while no Aβ aggregates were added to the control group. Meanwhile, we used solutions of ADLumin-5/Aβ aggregates as a positive control to reference the degraded bands. After electrophoresis, the gels are directly subjected to an image. As expected, we observed a band at the same position as the positive control from both chemiluminescence and fluorescence imaging (Fig.7a), suggesting that ADLumin-5 can degrade Aβ in a biologically relevant environment. To further confirm the capacity of ADLumin-5, we treated brain organoids that contain Aβs, and then used ELISA to analyze the concentration of Aβs. We found that ADLumin-5 could reduce Aβ burdens in organoids (Fig. 7b, SI Fig. 8). Our solution-based assays indicated that ADLumin-5 could degrade Aβ peptides, and TEM images showed thinner fibrils in the presence of ADLumin-5. To investigate whether ADLumin-5 alters the properties of Aβ plaques in tissues, we incubated ADLumin-5 with AD mouse brain tissue for 48 hours; the control group was similarly treated without ADLumin-5. Before and after treatment, the slides were stained with CRANAD-3, a near-infrared fluorescence probe for Aβ imaging. We quantified the fluorescence intensity of CRANAD-3 in each plaque and normalized it to the pre-treatment intensity. We found that both plaque core and peripheral decreased (SI Fig. 9a-c). To further confirm this effect, we first used tissue expansion technology to treat the tissue ^52^, and then treated the expanded tissue with ADLumin-5. As expected, we found that the intensity of the plaque core was slightly increased for the control group, while the intensity decreased 50% for the ADLumin-5 treated group (Fig. 7c-e), while the plaque shapes did not have significant alterations. Taken together, all the results suggested that ADLumin-5 could degrade Aβ peptides in a biologically relevant environment.

**Figure 7.**
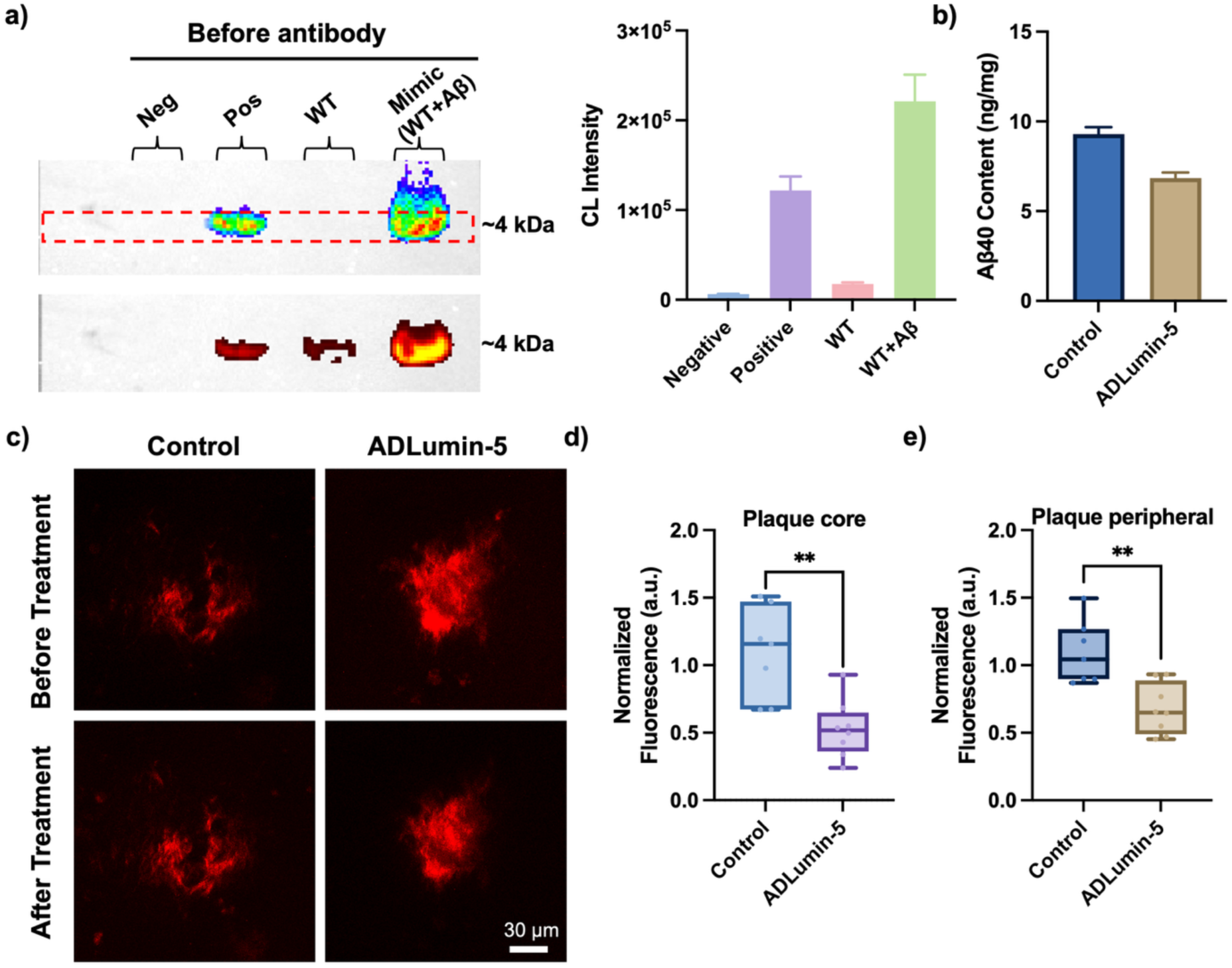
**a)** Brain homogenate with different treatments and degradation band quantification; **b**) ELISA analysis of Aβ concentration detection in 3D brain organoids after treatment with ADLumin-5; **c**) Study of Aβ degradation in fixed mouse brain tissue treated or untreated with ADLumin-5 (expanded sample); **d**) Normalized fluorescence intensity of plaque cores after treatment. Data are mean ± SD, n = 3. **p < 0.01; **e**) Normalized fluorescence intensity of plaque periphery after treatment. Data are mean ± SD, n = 3. **p < 0.01

### In vivo therapeutic studies and imaging monitoring

Our results from in vitro studies suggested that ADLumin-5 could have a potential therapeutic effect. In this regard, we performed longitudinal treatment with ADLumin-5 for 4 months. Meanwhile, we also used ADLumin-5 as an imaging probe to longitudinally monitor the changes in Aβ burdens. The imaging and treatment schedule is shown in Fig. 8a. We used three groups of mice for our study (WT group, ADLumin-5-treated group, and saline-treated group). The WT mice were used as a reference, as the imaging signals do not significantly change during the treatment period. The experimental 5xFAD group mice were treated with ADLumin-5 three times per week, and no ADLumin-5 treatment was given for 2 days before in vivo imaging. The control 5xFAD group was treated with a saline vehicle. At the beginning of treatment, we confirmed that ADLumin-5 could report the difference between 5xFAD mice and WT mice, as 2-fold higher signals were observed from the 5xFAD mice (Fig. 8b). Figure 8c revealed distinct Aβ levels among the groups. The vehicle-treated transgenic mice exhibited a gradual increase in Aβ concentrations over time, consistent with progressive amyloid accumulation in the disease model. In contrast, mice receiving the treatment showed a markedly attenuated increase, with Aβ levels remaining relatively stable throughout the study and even displaying a slight downward trend at later time points. This divergence between groups became more pronounced as the experiment progressed, indicating a sustained therapeutic effect. These results suggest that the treatment not only suppresses the accumulation of Aβ but may also facilitate its clearance or inhibit further aggregation. After the final treatment, the mice were sacrificed, and their brains were collected. Endpoint ELISA analysis confirmed reduced amyloid levels in treated animals, with decreases observed in both soluble and insoluble Aβ species compared to controls (Fig. 8d,e). We also conducted histological staining with brain slices using CRANAD-3, a well-characterized NIR fluorescent probe from our group ^53–56^. We found that plaque densities and plaque sizes from the ADLumin-5 treated group were lower and smaller, compared to the vehicle group, further supporting our in vivo imaging results (Fig. 8f, g and SI Fig.10). Together, these results demonstrate that ADLumin-5 enables simultaneous longitudinal imaging and therapeutic modulation of amyloid pathology in vivo, supporting its potential as a light-independent theranostic strategy.

**Figure 8.**
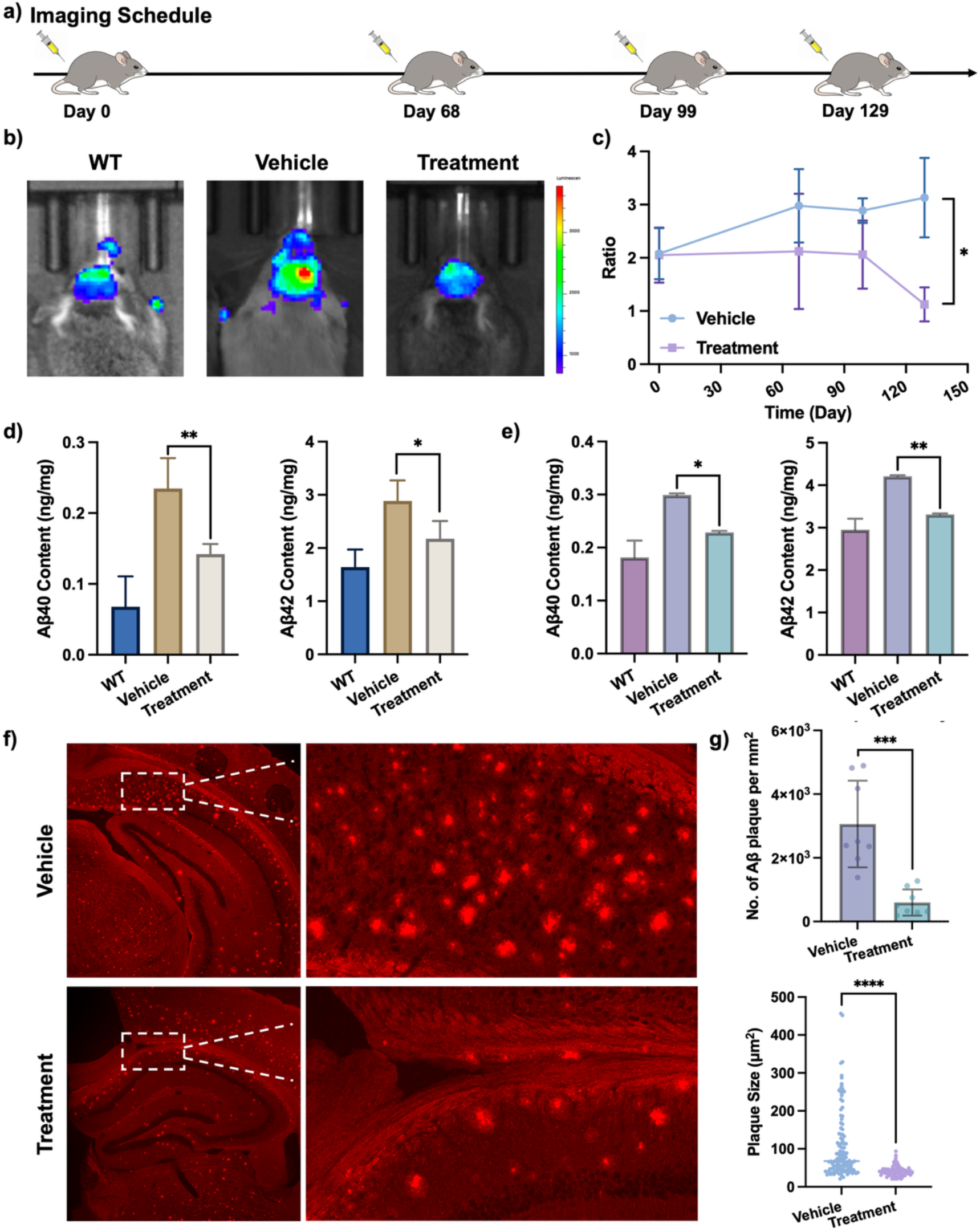
**a)** Treatment and imaging schedule; **b)** Representative in vivo images showing Aβ-associated signals in different treated mice; **c)** Quantification of the ratio chemiluminescent signals in the mouse brain over time. Data are mean ± SD, n = 5. *p < 0.05. **d)** ELISA analysis of the mouse brain soluble Aβ levels after 4 months of treatment. Data are mean ± SD, n = 3. *p < 0.05, **p < 0.01; **e**) ELISA analysis of the mouse brain insoluble Aβ levels after 4 months of treatment. Data are mean ± SD, n = 3. *p < 0.05, **p < 0.01. **f**) Ex vivo fluorescence imaging of Aβ burden in mouse brain sections with vehicle or ADLumin-5 treatment. **g**) Quantitative analysis of plaque density and plaque size in mouse brain sections with vehicle or ADLumin-5 treatment.

## Discussion

### Molecular light is adequate for photoreactions in vitro and in vivo

Phototherapy has been widely investigated and applied for a variety of diseases; however, the penetration limitation of light has severely hampered its applications in deep targets. Therefore, it is highly desirable if the tissue penetration limitation of light can be overcome. Recent studies from our group and other groups have suggested that molecularly produced light (molecular light) could be a viable approach to partially overcome the penetration problem ^15,25,26,57^. In conventional phototherapy, a large photon flux can be shone toward the target area; however, only a small fraction of these photons can reach deep tissue targets, as the majority of the photons are absorbed by non-targeted tissues, scattered away before reaching the target, and reflected by the surface ^58^. In contrast, while the total photon flux from molecular light is significantly lower, it can be delivered to the target at zero to nanometer distances. This close proximity significantly enables effective energy transfer through absorption as well as chemo- or bioluminescence resonance energy transfer. Approximately, molecular light could bring the light source ∼ 10^6^ times closer to the target, compared to the externally shone light (nanometers versus centimeters). Furthermore, due to the principles of quantum mechanics, each photon emitted by molecular light at such close proximity could be absorbed by the target if its absorbance wavelength aligns. These factors suggest that molecular light may have an equivalent potential to initiate photoreactions in vitro and in vivo. In this study, again, we demonstrated that molecular light is adequately efficient to initiate photo-reactions in vitro and in vivo with misfolded proteins in mouse models.

### Mechanism of photo-oxidation and photo-degradation

The limited results from our study here it indicates that higher light-emitting efficiency, higher degradation, and oxidation. However, it is not clear what the major facts are that govern the reactions. In Kanai’s report, they used a heavy atom to increase the production of ^1^O_2_ via ISC of the chemi-excited intermediates ^26^. In their study, they proposed a possible mechanism for the oxidation of methionine-35 in Aβ peptides. One is the HAT mechanism that abstracts one hydrogen of the methionine side chain and transfers it to the IPO moiety. Interestingly, they provide experimental data to support that the ROS species is superoxide anion, instead of ^1^O_2_. These results are also different from our studies, in which we showed that ADLumin-5 can induce the generation of O_2_^-.^ and ^1^O_2_. This might be the reason that no halogen is needed in the ADLumin-X to execute its photo-oxidation/degradation capacity. More theoretical studies are needed and are being conducted in our group.

### Self-photosensitizing efficiency of ADLumin-5

We observed ∼6% ^1^O_2_ production within 60 minutes, which is relatively low compared with methylene blue (MB) under LED irradiation. However, over extended periods (e.g., 24 hours or longer), the cumulative ^1^O_2_ generation increased substantially (∼70%). This moderate efficiency may be advantageous, as high levels of ^1^O_2_ are difficult to control and can be highly cytotoxic, whereas slower, sustained production may be more controllable and associated with reduced toxicity. As this is the first report of self-photosensitizing activity for this compound, further studies are needed to determine how controlled ^1^O_2_ generation influences its degradation/photo-oxidation capacity while maintaining minimal toxicity.

### Photodegradation/photo-fragmentation

Protein degradation has recently been intensively and comprehensively investigated for seeking therapies and understanding the fundamental mechanisms of different diseases. PROTAC, Lyo-TAC, and others have been demonstrated to efficiently degrade the “bad” proteins. PDT therapy has been comprehensively and intensively studied for cancer research and treatment and other diseases; however, it is rarely investigated whether PDT can lead to protein degradation. In this work, we showed that DEMOL could be used to cleave misfolded proteins. We believe it is highly likely that PDT and DEMOL can cause protein degradation in cancer tissue. Indeed, it has been reported that photo-oxidation can cause fragmentation of proteins ^59–62^, and recent photo-oxidative reactions have been shown to facilitate bond cleavage in proteins ^62^. However, it will be interesting to investigate whether and how “bad” proteins can be degraded during PDT and DEMOL. In this study, we demonstrated that molecular light can be self-photosensitizing and cause protein oxidation and cleavage, particularly the oxidation and degradation of methionine. Photo-oxidation of proteins can be a double-edged sword. For a protein with normal function, its photo-oxidation is harmful; in contrast, photodynamic therapy uses this oxidation reaction to damage/degrade the “bad” proteins that are pathologically related to cancers, infections, and others ^63,64^. In neurodegenerative diseases, misfolded proteins, such as Aβs and tau, are considered “bad” proteins ^38^. In this regard, photo-oxidation can be harnessed for seeking effective treatments for Alzheimer’s disease and other misfolded protein-related diseases. We believe that precisely controlling the degradation may elevate the precision of PDT and photo-therapy and increase its efficiency and/or reduce resistance. In addition, DEMOL with ADLumin-Xs may potentially be applied to degrade dysfunctional proteins, thereby helping maintain protein homeostasis by enhancing the clearance of aberrant proteins.

### Beta-sheet specificity with ADLumin-5 for photo-oxidation and degradation

Our model study with the PAM-K2 peptide strongly supported that ADLumin-5 induced photo-oxidation and degradation was highly beta-sheet specific. Notably, we also clearly demonstrated that ADLumin-5 was generic for photo-oxidation and photo-degradation of misfolded proteins, including not only Aβ aggregates, but also tau, alpha-synuclein, and TDP-43. Considering that most misfolded proteins consist of beta-sheet proteins/peptides, our study suggests that molecular light could be generally used to reduce the harmful effects of misfolded proteins via increasing polarity and fragmentation of the proteins. Regarding ADLumin-X’s specificity to beta-sheet aggregation, it is likely due to its binding to the hydrophobic groove, since our previous study indicated that ADLumin-X can be docked into the hydrophobic grooves of a variety of fibrils of misfolded proteins ^36^. However, better specificity can be achieved if we know the exact binding sites. With this knowledge, it will greatly assist in rationally designing ADLumin-X for degradation and oxidation. However, such precise binding has not been validated with Cryo-EM, and our group is currently working in this direction.

### Sites and consequences of photoreactions

Our results indicate that the primary site of photo-induced reactions in amyloid-β (amyloid beta) is centered around Met35 (M35). This residue appears to be involved not only in oxidation but also in cleavage (degradation) and conjugation processes. This finding is both interesting and significant, as oxidation at M35 can disrupt the hydrophobic core of Aβ peptides, thereby destabilizing fibrillar structures and facilitating their disassembly or clearance.

The region encompassing residues 25–35 (Aβ25–35) is widely recognized as a central hydrophobic domain that retains much of the toxicity of the full-length peptide ^48,65,66^, and has been extensively used in experimental models of neurotoxicity and Alzheimer’s disease. In addition, recent work suggests that residues I32–M35–V40 form a triad that is energetically favorable for aggregation ^67^. Thus, photo-induced modification at M35 may disrupt a critical structural motif, potentially reducing aggregation propensity and toxicity. Consistent with prior reports, our data also indicate that oxidized Aβ species exhibit reduced toxicity. Importantly, our study shows that the photoreaction induces not only oxidation but also peptide cleavage and conjugation. To our knowledge, this is the first report demonstrating such conjugation products. While previous studies (e.g., Kanai and others) have shown that molecular light or LED-induced photoreactions can promote Aβ oxidation ^26,44–46,68^, they did not report evidence of peptide cleavage or conjugation. Although the term “degradation” has been used in several publications, direct evidence for peptide bond cleavage has generally been lacking ^69–71^.

For other misfolded proteins, such as α-synuclein and tau, cleavage may also occur at methionine residues. However, in sequences that lack methionine, photo-oxidation is more likely to occur at residues such as cysteine, tyrosine, or histidine. Oxidation at these sites can lead to peptide cleavage and generate fragments with the observed molecular weights, as these amino acids are also susceptible to photo-induced oxidation ^44,68,72^.

### In vivo phototheranostic studies

In this study, we evaluated the theranostic potential of ADLumin-5 in the 5xFAD mouse model as a proof of concept. Our in vitro results indicate that ADLumin-5 functions as a general chemiluminescent probe capable of binding to and photo-oxidizing a range of misfolded proteins. Accordingly, it is conceivable that ADLumin-5 could be applied as a theranostic agent in other neurodegenerative disease models. Ongoing studies in our laboratory are exploring these possibilities, and the results will be reported in due course. In this report, behavioral assessments of memory and recognition were not conducted, as the primary objective was to establish a proof of concept for phototherapy using self-photosensitizing chemiluminescent compounds. In addition, the sample size in each group (n = 3–5) is too small to support reliable behavioral analysis.

## Conclusion

In summary, this study demonstrates that ADLumin-5 can emit light while simultaneously exerting self-photosensitizing activity. This property initiates photoreactions that induce oxidation and degradation of a range of misfolded proteins, including amyloid beta, tau, α-synuclein, and TDP-43, as supported by SDS-PAGE, mass spectrometry, and Western blot analyses. Furthermore, photo-oxidized Aβ species exhibit altered biochemical properties, including reduced seeding capacity, increased susceptibility to proteolytic degradation, and changes in fibril size. Results from cellular and organoid models further demonstrate that ADLumin-5 attenuates Aβ-associated neurotoxicity. Finally, in vivo studies in the 5xFAD mouse model are consistent with the in vitro findings, showing that ADLumin-5 can function as a theranostic agent to both monitor treatment response and reduce Aβ accumulation. We believe that our approach provides a new avenue for therapeutic discovery in neurodegenerative diseases.

## Acknowledgments

This research has been supported by NIA grants R01AG083759 (C.R.), R01AG085562 (C.R.), R21AG080222 (C.R.), R21AG078749 (C.R.), and S10OD028609 (C.R.). We thank ACRO Biosystems for providing the brain organoids.

---

**Scheme 1.**
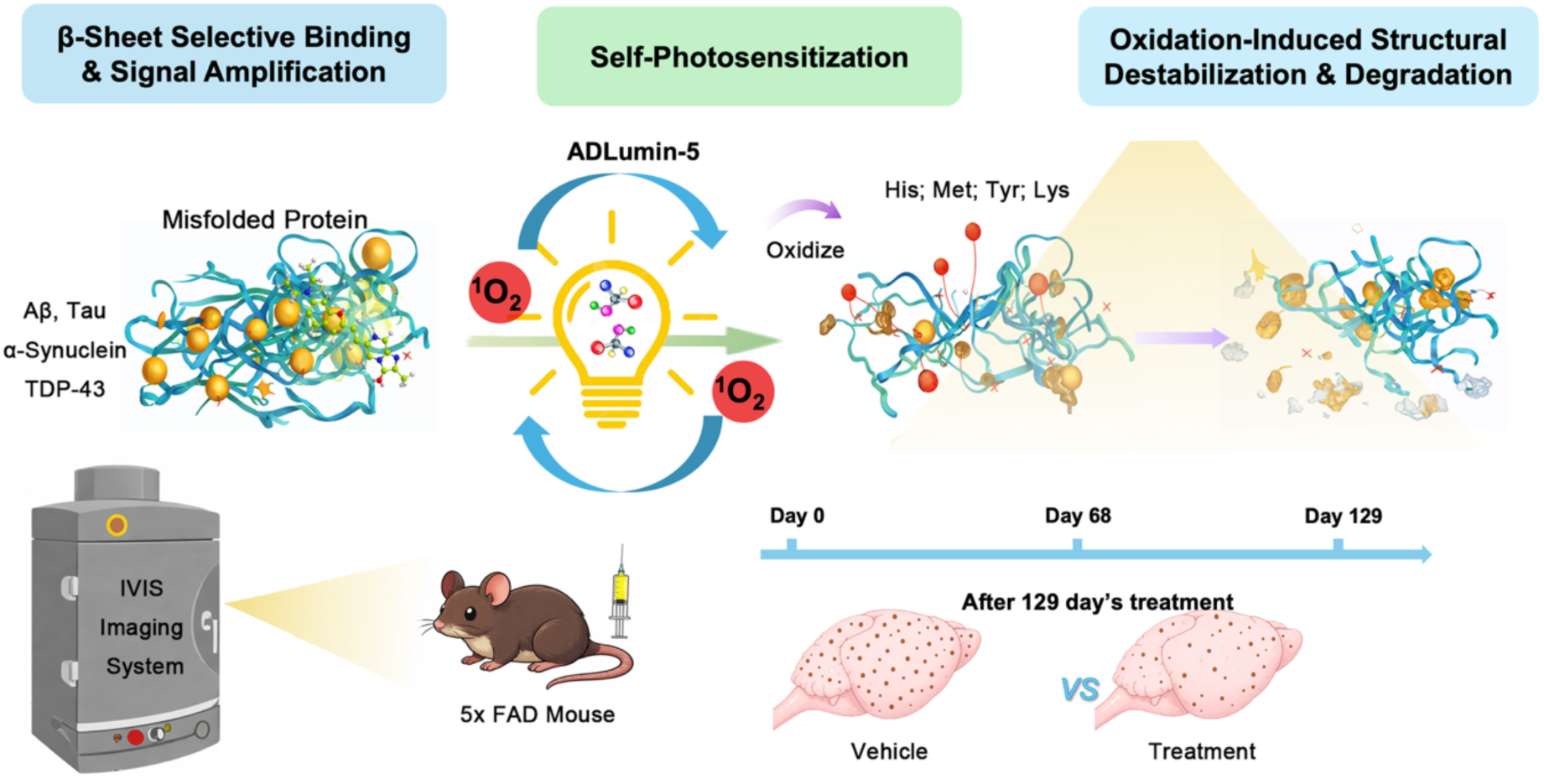
Schematic illustration of the mechanism of ADLumin-5-mediated clearance of amyloid-β (Aβ) aggregates. ADLumin-5 enables self-sustained generation of singlet oxygen (^1^O_2_) through molecular light emission, leading to oxidative modification and degradation of misfolded Aβ species.

