## Supplemental Information for "Targeted Photodegradation of Misfolded Proteins via Self-photosensitizing with Molecularly Produced Light"

**Molecularly Produced Light for Photooxidation and Photodegradation of Misfolded Proteins**

**Material and instrument**

Aβ_1–40_, Aβ_1-42_ peptide and Alpha-Synuclein Preformed Fibrils was purchased from rPeptide, and different forms of Aβ_1–40_, Aβ_1-42_, including the monomer, oligomer and aggregate were prepared by our previously reported methods. ^[1]^ Human Recombinant Tau-441 (2N4R) P301S Mutant Protein Filaments was purchased from Stressmarq. Human TDP-43 protein (His Tag) was obtained from Acrobiosystems. Aβ_1-42_ (FAM-labeled) was purchased from Anaspec. Palmitic-VVVMAAAKKK (PAM-K2) was customized in Genescript. Purified anti-β-Amyloid 1-16 (clone 6E10) was purchased from BioLegend, Tau Monoclonal Antibody (TAU-5) and TDP-43 Recombinant Rabbit Monoclonal Antibody (JM51-10) were obtained from Invitrogen, Anti-Alpha-synuclein antibody (MJFR1) was purchased from Abcam. Goat anti Rabbit IgG (H+L) secondary antibody (HRP), Goat anti Mouse IgG (H+L) secondary antibody (HRP) was purchased from Thermo Fisher Scientific. UV spectrum was performed on SpectraMax Plus 384 Microplate Reader (Molecular Devices). Fluorescence measurements were carried out using an F-7100 fluorescence spectrophotometer (Hitachi). Liquid chromatography-mass spectrometry (LC-MS) was performed using an Agilent 1200 Series apparatus with an LC/MSD trap and Daly conversion dynode detector with UV detection at 254 nm. Electrophoresis Power Supply-EPS 3500 (Pharmacia) and XCell SureLock™ Mini-Cell Tank (Invitrogen™) were used for SDS-PAGE and Western Blot. Wild-type (B6SJLF1/J) female mice and APP female mice (B6; C3-Tg (APPswe, PSEN1dE9)85Dbo/Mmjax) were purchased from Jackson Laboratory. All animal experiments were approved by the Institutional Animal Use and Care Committee at Massachusetts General Hospital. Animals were housed in groups in standardized cages with a 12/12 h light/dark cycle with unrestricted access to food and water at room temperature (25°C) with 55% humidity. The IVIS Spectrum imaging system (PerkinElmer) was used for in vitro and in vivo imaging.

**Experimental section**

**Preparation of Aβ_40_ aggregates**.

1.0 mg of Aβ_40_ peptide (TFA) was suspended in a 0.05 % ammonia hydroxyl solution (1.0 mL), then 200.0 μL of the resulting solution was taken and diluted 5-fold with PBS buffer (pH 7.4) and stirred at room temperature for 3 days to obtain 50 μM of Aβ_40_ aggregates. Transmission electron microscopy and fluorescence tests with Thioflavin T were used to confirm the formation of aggregates.

**Preparation of Aβ_42_ aggregates.**

1.0 mg of Aβ_42_ peptide (TFA) was suspended in a 0.15 % ammonia hydroxyl solution (1.0 mL), then 200.0 μL of the resulting solution was taken and diluted 5-fold with PBS buffer (pH 7.4) and stirred at room temperature for 3 days to obtain 50 μM of Aβ_40_ aggregates. Transmission electron microscopy and fluorescence tests with Thioflavin T were used to confirm the formation of aggregates.

**Preparation of FAM-Aβ_42_ aggregates.**

1.0 mg of FAM-Aβ_42_ peptide was suspended in 1.0 mL PBS, then 250.0 μL of the resulting solution was taken and diluted 4-fold with PBS buffer (pH 7.4) and stirred at room temperature for 8 hours to obtain 50 μM of FAM-Aβ_42_ aggregates. Transmission electron microscopy and fluorescence tests with Thioflavin T were used to confirm the formation of aggregates.

**Preparation of PAM-K2 Aggregates**.

1.0 mg of PAM-K2 peptide was dissolved in 1.56 mL PBS buffer (1X, pH = 7.4) and incubated for 72 h with constant shaking to obtain 500 μM of PAM-K2 aggregates.

**In vitro spectral test in solutions.**

ADLumin-5 solutions (50 μM) in 3.0 mL DMSO were prepared, and UV spectra of the prepared solutions were recorded using SpectraMax Plus 384 Microplate Reader (Molecular Devices). ADLumin-5 in 1.0 mL of DMSO (250.0 nM), PBS (10 μM), and DCM (100 nM) were prepared, and fluorescence spectra of the prepared solutions were recorded using an F-7100 fluorescence spectrophotometer (Hitachi). A blank control of DMSO was used to correct the final spectra.

PAM-K2 monomer or PAM-K2 aggregate (5 μM) in PBS solutions (1.0 mL) containing Thioflavin T (0.5 μM, 1% DMSO), fluorescence spectra of the prepared solutions were recorded.

Aβ40/42 aggregate (final concentration: 10 μM) and ADLumin-5 (final concentration: 10 μM, 1% DMSO) were dissolved in PBS solutions (1.0 mL) and the fluorescence spectra of the prepared solutions were recorded.

**Singlet oxygen (^1^O_2_) detection by ADPA.**

1) The solution of ADLumin-5 (25 μM, 1% DMSO) in PBS solutions (3.0 mL) containing ADPA (25 μM, 1% DMSO).

2) The solution of MB (25 μM, 1% DMSO) in PBS solutions (3.0 mL) containing ADPA (25 μM, 1% DMSO).

3) The solution of ADLumin-5 (25 μM, 1% DMSO), MB (25 μM, 1% DMSO) in PBS solutions (3.0 mL) containing ADPA (25 μM, 1% DMSO).

4) A solution containing ADPA (25 μM, 1% DMSO) only was used as the control. The absorbance was measured every 15 seconds, with continuous monitoring for 1 hour in dark conditions.

5) The solution of ADLumin-1 (10 μM, 1% DMSO) in PBS solutions (3.0 mL) containing ADPA (10 μM, 1% DMSO).

6) Different concentrations of ADLumin-5 (1, 10, 50, 100 μM, 1% DMSO) in PBS solutions (3.0 mL) containing different concentrations of ADPA (1, 10, 50, 100 μM, 1% DMSO).

7) The solution of ADLumin-15 (10 μM, 1% DMSO) in PBS solutions (3.0 mL) containing ADPA (10 μM, 1% DMSO).

8) The solution of ADLumin-5 (10 μM, 1% DMSO) in PBS solutions (3.0 mL) containing Aβ40 aggregate (10 μM) and ADPA (10 μM, 1% DMSO).

9) The solution of ADLumin-5 (10 μM, 1% DMSO) in PBS solutions (3.0 mL) containing Aβ42 aggregate (10 μM) and ADPA (10 μM, 1% DMSO). The absorbance was measured every 30 seconds, with continuous monitoring for 240 min in dark conditions.

10) The solution of ADLumin-Xs (X=1, 4, 5, 6, 9, 10, 15) (100 μM, 1% DMSO) in PBS solutions (3.0 mL) containing ADPA (100 μM, 1% DMSO). The absorbance was measured every 2 min after LED (THORLABS (LIU470A); 470 nm LED Array; 24 V; 15 W) irradiation.

**Singlet oxygen (^1^O_2_) Measurement with FFA.**

The solution of ADLumin-Xs (X=1, 4, 5, 6, 9, 10, 15) (100 μM, 1% DMSO) in PBS solutions (1.0 mL) containing ADPA (100 μM, 1% DMSO).

To a solution of Aβ_40_ or Aβ_42_ aggregate (final concentration 10 μM) was added ADLumin-5 (final concentration: 10 μM) and furfuryl alcohol (final concentration: 100 μM). The reaction mixture was incubated at 37 ℃ for 24 h. The remaining furfuryl alcohol was quantified by UV absorbance using high-performance liquid chromatography (HPLC).

**Superoxide anion radical (O_2_^•−^) detection by DHE.**

Evaluation of O_2_^•−^ was performed by the interaction between dihydroethidine and DNA.^[2]^ Dihydroethidium (DHE) was utilized as O_2_^•−^ specific probe because it could intercalate in DNA and emit red fluorescence upon interacting with O_2_^•−^. Briefly, ADLumin-Xs (X=1, 4, 5, 6, 9, 10, 15) (25 μM, 1% DMSO) and DHE (25 μM, 1% DMSO) were dissolved in PBS solutions (3.0 mL) containing ctDNA (250.0 μg/mL). A mixture containing DHE and ctDNA only was used as the control. The fluorescence spectra were measured every 30 seconds.

**Catechol study.**

Aβ_40_ or Aβ_42_ aggregate (final concentration: 2 μM) and ADLumin-5 (final concentration: 2 μM, 1% DMSO) were dissolved in PBS solutions (1.0 mL) and incubated for 24 h, then different concentration of Catechol (0 eq, 5 eq, and 10 eq) was separately added to the above solution. Measure the fluorescence changes for different time points.

**In vitro chemiluminescence spectral test in solutions.**

ADLumin-Xs (X=1, 4, 5, 6, 9, 10, 15) (25 μM, 1% DMSO) were prepared, chemiluminescence intensity of the prepared solutions were seeded into 96-well plate and recorded with the IVIS spectrum imaging system (PerkinElmer). A blank control of 10% DMSO/PBS was used to correct the final spectra.

PAM-K2 monomer (50 μM) or PAM-K2 aggregate (50 μM) in PBS solutions (1.0 mL) containing ADLumin-5 (50 μM, 1% DMSO), chemiluminescence intensity of the prepared solutions was recorded.

Aβ_40_ monomer/aggregate, Aβ_42_ monomer/aggregate, alpha-synuclein monomer/aggregate, Tau-441 2N4R monomer/aggregate, and TDP-43 monomer/aggregate (final concentration: 2 μM) were separately added with ADLumin-5 (final concentration: 2 μM, 1% DMSO), chemiluminescence intensity of the prepared solutions was seeded into a 96-well plate and recorded with the IVIS spectrum imaging system for 200 min.

**Recovery study.**

Aβ_40_ or Aβ_42_ aggregate (final concentration: 2 μM) and ADLumin-1/-5/-15 (final concentration: 2 μM, 1% DMSO) were dissolved in PBS solutions (1.0 mL).

Treated group: The emission intensity was recorded.

Treated for 24 h: After incubating the mixture for 24 h, the emission intensity was recorded.

Treated for 24 h + fresh: After incubating the mixture for 24 h, fresh ADLumin-1/-5/-15 was added to the samples, and the emission intensity was recorded.

Treated for 48 h: After incubating the mixture for 48 h, the emission intensity was recorded.

Treated for 48 h + fresh: After incubating the mixture for 48 h, fresh ADLumin-1/-5/-15 was added to the samples, and the emission intensity was recorded.

**Protease K digest experiments.**

Six groups of Aβ_40_ or Aβ_42_ aggregates (final concentration: 2 μM) in PBS (1.0 mL) were treated as follows, respectively.

Group 1: The Aβ_40_ or Aβ_42_ aggregates were incubated for 24 h, then mixed with ADLumin-1 (final concentration: 2 μM, 1% DMSO).

Group 2: The Aβ_40_ or Aβ_42_ aggregates incubated with ADLumin-1(final concentration: 2 μM, 1% DMSO) for 24 h.

Group 3: The Aβ_40_ or Aβ_42_ aggregates were incubated for 24 h, then mixed with ADLumin-5 (final concentration: 2 μM, 1% DMSO).

Group 4: Aβ_40_ or Aβ_42_ aggregates incubated with ADLumin-5(final concentration: 2 μM, 1% DMSO) for 24 h.

Group 5: The Aβ_40_ or Aβ_42_ aggregates were incubated for 24 h, then mixed with ADLumin-15 (final concentration: 2 μM, 1% DMSO).

Group 6: The Aβ_40_ or Aβ_42_ aggregates incubated with ADLumin-15 (final concentration: 2 μM, 1% DMSO) for 24 h.

After these treatments, protease K (final concentration: 0.1 mg/mL) was added to each sample, and all these samples were incubated in a 37 ℃ humidity incubator, and the data were recorded every 30 min.

**Seeding experiments.**

The Aβ_40_ or Aβ_42_ aggregates (300 μL, 25 μM) in PBS were treated as follows:

Group 1: Aβ_40_ or Aβ_42_ aggregates were kept in a dark environment for 24 h.

Group 2: Aβ_40_ or Aβ_42_ aggregates were mixed with ADLumin-5 (1 eq), and the resulting mixture was kept in a dark environment for 24 h.

After the treatments, all these samples were added to a solution of Aβ_40_ monomers (6 mL, 25 μM) separately. At every time-point, a 500 μL sample was pipetted out. Then the emission intensity was recorded at 450 nm (Ex = 420 nm). The emission intensities were recorded twice daily until no significant intensity change could be observed. The emission peak intensities were normalized by the last data point for each group.

**Seletivity Study.**

Aβ_40_ monomer/aggregates, Aβ_42_ monomer/aggregates and non-amyloid off-target model peptides containing oxidation-sensitive amino acids residues, angiotensin IV (Ang IV), neurokinin A (NKA), and RNase A (4 μM) was separately treated with ADLumin-5, fluorescence intensity of the prepared solutions were seeded into 96-well plate and recorded with the IVIS Spectrum imaging system (Ex = 460 nm; Em = 520 nm).

**Size and Zeta Potential.**

Aβ_40_ or Aβ_42_ aggregate (final concentration: 400 nM) incubated with or without ADLumin-5 (final concentration: 400 nM) at 37 ℃ for 24 h, and record the size and zeta potential changes through Dynamic light scattering (DLS).

**Oxygenation assay of PAM-K2.**

To a solution of PAM-K2 monomer/aggregate (final concentration: 20 μM) in PBS (100 μL), ADLumin-Xs (X=1, 4, 5, 6, 9, 10, 15) (20 μM, 1% DMSO) was added, and the solution was incubated in the dark at 37 °C for 24 h. Then, LC-MS was applied to analyze the oxygenation yield and cleavage. For each fragment of interest, the oxygenation yield was calculated as follows: oxygenation yield [%] = (MS area of oxygenated fragment) / (sum of MS areas of intact and oxygenated fragment) × 100.

**Oxygenation assay of Aβ and off-target peptides.**

To a solution of aggregated Aβ and off-target peptide Angiotensin IV (20 μM) in PBS (100 μL), ADLumin-5 (20 μM, 1% DMSO) was added, and the solution was incubated in the dark at 37 °C for 24 h. The mixture was analyzed using MALDI-TOF MS after desalting with ZipTip U-C18. The oxygenation yield was calculated as follows: oxygenation yield [%] = (sum of MS intensities of oxygenated species) / (sum of MS intensities of remaining and oxygenated species) × 100.

**Oxygenation and Cleavage Study of protein with β-sheets**.

To a solution of Aβ_40_ or Aβ_42_ aggregate (final concentration: 20 μM) in PBS (100 μL), ADLumin-5 (final concentration: 20 μM) was added, and the mixture was incubated in the dark at 37 °C for 24 h. Then the mixture was subjected to enzymatic digestion with Trypsin Gold (Promega Co.) (1 μL, 1 μg/μL in 50 mM acetic acid aqueous solution) and Protease Max (Promega Co.) (0.25 μL, 10 μg/μL in 50 mM ammonium bicarbonate aqueous solution) at 37 °C for 12-18 h. and then analyzed using LC-MS.

To a solution of αSyn aggregate (final concentration: 20 μM) in PBS (100 μL), ADLumin-5 (final concentration: 20 μM) was added, and the mixture was incubated in the dark at 37 °C for 24 h. Then the mixture was subjected to enzymatic digestion with Trypsin Gold (Promega Co.) (1 μL, 1 μg/μL in 50 mM acetic acid aqueous solution) and Protease Max (Promega Co.) (0.25 μL, 10 μg/μL in 50 mM ammonium bicarbonate aqueous solution) at 37 °C for 12-18 h. and then analyzed using LC-MS.

To a solution of tau-441 2N4R (final concentration: 20 μM) in PBS (100 μL), ADLumin-5 (final concentration: 20 μM) was added, and the mixture was incubated in the dark at 37 °C for 24 h. The mixture was subjected to sonication after the addition of Tris-HCl buffer (1900 μL, 50mM, pH 6.8) with urea (8 M) and DTT (10 mM), and the solvent was exchanged to ammonium bicarbonate aqueous solution (400 μL, 50 mM) with Amicon Ultra (30 kDa, Merck Millipore Ltd.). Subsequently, oxygenated tau was digested with Trypsin Gold (1 μL, 1 μg/μL in 50 mM acetic acid aqueous solution) and Protease Max (1 μL, 10 μg/μL in 50 mM ammonium bicarbonate aqueous solution) at 37 °C for 12-18 h. After evaporating the solvent, the pellet was dissolved with aqueous formic acid (80 μL, 0.1% (v/v)), and then the supernatant after centrifugation (21,300 g, 10 min, 4 °C) was analyzed using LC-MS/MS.

**Identification of the oxygenation position of Aβ.**

Aβ_40_ aggregates (final concentration: 20 μM) were incubated with ADLumin-5 (final concentration: 20 μM; 1% DMSO) for 24 hours. After incubation, 10 μL of the mixture was loaded per well onto a 4–20% SDS-PAGE gel. Electrophoresis was performed at 80 V for 20 minutes, followed by 120 V for 35 minutes. After electrophoresis, the gel was stained with Coomassie Brilliant Blue. The band of interest was excised using a sterile scalpel, then cut into 2 × 2 mm cubes and transferred into a 1.5 mL snap-cap microcentrifuge tube.

To destain the gel pieces, 75 μL of a 1:1 solution of 100 mM ammonium bicarbonate and acetonitrile was added. The samples were briefly vortexed and incubated at room temperature for 15 minutes. The solution was discarded, and this wash step was repeated once more. After the second wash, 75 μL of 100% acetonitrile was added. Once the gel pieces shrank and turned opaque white (~30 seconds to 1 minute), the acetonitrile was removed.

Gel pieces were rehydrated with 75 μL of 10 mM DTT in 50 mM ammonium bicarbonate and incubated at 56 °C for 1 hour in a water bath. Following incubation, the tubes were briefly centrifuged, and the DTT solution was discarded. Then, 75 μL of 55 mM iodoacetamide in 50 mM ammonium bicarbonate was added, and the samples were incubated in the dark at room temperature for 30 minutes. The iodoacetamide solution was removed.

Next, gel plugs were washed twice with 75 μL of a 1:1 solution of acetonitrile and 100 mM ammonium bicarbonate to remove residual Coomassie dye. If necessary, additional washes were performed until the dye was fully removed. Finally, 75 μL of 100% acetonitrile was added; after the gel pieces shrank and turned opaque white (~30 seconds to 1 minute), the acetonitrile was removed.

For digestion, gel pieces were rehydrated with digestion buffer (50 mM ammonium bicarbonate, 5 mM CaCl₂, and 5 ng/μL trypsin) at 4 °C. A sufficient volume was added to cover the gel pieces. After 15 minutes on ice, the buffer was removed and replaced with 70 μL of 50 mM ammonium bicarbonate containing 5 mM CaCl₂. Digestion was carried out at 37 °C overnight in a warm-air incubator.

The following day, tubes were briefly centrifuged, and the supernatant was collected into new, labeled 1.5 mL tubes. The remaining gel pieces were sequentially extracted with ~60 μL of 50% acetonitrile, 0.3% formic acid for 15 minutes, followed by 80% acetonitrile, 0.3% formic acid for another 15 minutes. Each supernatant was pooled with the previous extracts.

The combined extracts were frozen at –80 °C for 30 minutes, then dried using a speed vacuum concentrator. Samples were then desalted using zip-tip or stage-tip protocols and subjected to LC-MS analysis.

**SDS-PAGE Electrophoresis.**

PAM-K2 aggregates (final concentration: 20 μM) were incubated with ADLumin-5 (1 eq or 2.5 eq; 1% DMSO) for 24 hours. After incubation, 10 μL of the mixture was loaded per well onto a 4–20% SDS-PAGE gel. Electrophoresis was performed at 80 V for 20 minutes, followed by 120 V for 30 minutes. After electrophoresis, cut off the gel and record with the IVIS spectrum imaging system (PerkinElmer).

FAM-Aβ_40_/ FAM-Aβ_42_ aggregates (final concentration: 20 μM) were incubated with ADLumin-1/5/15 (final concentration: 20 μM; 1% DMSO) separately for 24 hours. After incubation, 10 μL of the mixture was loaded per well onto a 4–20% SDS-PAGE gel. Electrophoresis was performed at 80 V for 20 minutes, followed by 120 V for 30-40 minutes. After electrophoresis, cut off the gel and record with the IVIS spectrum imaging system (PerkinElmer).

**Western Blot Assay.**

Aβ_40_/Aβ_42_/Alpha-synuclein/Tau-441 2N4R/TDP-43 aggregates (final concentration: 20 μM) were incubated with ADLumin-1/5/15 (final concentration: 20 μM; 1% DMSO) separately for 24 hours. After incubation, 10 μL of the mixture was loaded per well onto a 4–20% SDS-PAGE gel. Electrophoresis was performed at 80 V for 20 minutes, followed by 120 V for 30-40 minutes. Equilibrate gel and membrane (PVDF or nitrocellulose) in transfer buffer. Activate PVDF in methanol before use. Assemble the transfer sandwich then transfer at 35 V, 100 mA for 45 minutes at 4°C. After the transfer, the PVDF membrane was imaged in the IVIS fluorescence channel and chemiluminescence channel, respectively. Subsequently, the samples were blocked with 5% BSA and incubated with 6E10 (1:1000, Biolegend) at 4°C overnight. After incubation with HRP-conjugated goat anti-mouse secondary antibody (1:2000, Thermo Fisher Scientific), the protein was analyzed by ECL reagent and quantified by the IVIS system.

Alpha-synuclein aggregates (final concentration: 20 μM) were incubated with ADLumin-1/5/15 (final concentration: 20 μM; 1% DMSO) separately for 24 hours. After incubation, 10 μL of the mixture was loaded per well onto a 4–20% SDS-PAGE gel. Electrophoresis was performed at 80 V for 20 minutes, followed by 120 V for 45 minutes. Equilibrate gel and membrane (PVDF or nitrocellulose) in transfer buffer. Activate PVDF in methanol before use. Assemble the transfer sandwich then transfer at 35 V, 100 mA for 45 minutes at 4°C. After the transfer, the PVDF membrane was imaged in the IVIS fluorescence channel and chemiluminescence channel, respectively. Subsequently, the samples were blocked with 5% BSA and incubated with anti-alpha-synuclein antibody RabMab (1:1000, Abcam) at 4°C overnight. After incubation with HRP-conjugated goat anti-rabbit secondary antibody (1:2000, Thermo Fisher Scientific), the protein was analyzed by ECL reagent and quantified by the IVIS system.

Human Recombinant Tau-441 (2N4R) P301S Mutant Protein Filaments (final concentration: 20 μM) were incubated with ADLumin-1/5/15 (final concentration: 20 μM; 1% DMSO) separately for 24 hours. After incubation, 10 μL of the mixture was loaded per well onto a 4–20% SDS-PAGE gel. Electrophoresis was performed at 80 V for 20 minutes, followed by 120 V for 60 minutes. Equilibrate gel and membrane (PVDF or nitrocellulose) in transfer buffer. Activate PVDF in methanol before use. Assemble the transfer sandwich then transfer at 35 V, 100 mA for 80 minutes at 4°C. After the transfer, the PVDF membrane was imaged in the IVIS fluorescence channel and chemiluminescence channel, respectively. Subsequently, the samples were blocked with 5% BSA and incubated with Tau Monoclonal Antibody (TAU-5,1:500, Invitrogen) at 4°C overnight. After incubation with HRP-conjugated goat anti-mouse secondary antibody (1:2000, Thermo Fisher Scientific), the protein was analyzed by ECL reagent and quantified by IVIS system.

TDP-43 aggregates (final concentration: 20 μM) were incubated with ADLumin-1/5/15 (final concentration: 20 μM; 1% DMSO) separately for 24 hours. After incubation, 10 μL of the mixture was loaded per well onto a 4–20% SDS-PAGE gel. Electrophoresis was performed at 80 V for 20 minutes, followed by 120 V for 60 minutes. Equilibrate gel and membrane (PVDF or nitrocellulose) in transfer buffer. Activate PVDF in methanol before use. Assemble the transfer sandwich then transfer at 35 V, 100 mA for 80 minutes at 4°C. After the transfer, the PVDF membrane was imaged in the IVIS fluorescence channel and chemiluminescence channel, respectively. Subsequently, the samples were blocked with 5% BSA and incubated with TDP-43 Recombinant Rabbit Monoclonal Antibody (JM51-10, 1:1000, Invitrogen) at 4°C overnight. After incubation with HRP-conjugated goat anti-rabbit secondary antibody (1:2000, Thermo Fisher Scientific), the protein was analyzed by ECL reagent and quantified by the IVIS system.

For the time-dependent study, Aβ_40_ or Aβ_42_ aggregates (final concentration: 20 μM) were incubated with ADLumin-5 (final concentration: 20 μM; 1% DMSO) for different times, then followed the previous procedure.

For the concentration-dependent study, Aβ_40_ or Aβ_42_ aggregates (final concentration: 20 μM) were incubated with different concentrations of ADLumin-5 (1% DMSO) for 24 h, then followed the previous procedure.

**Cytotoxicity Assay**

SH-SY5Y cells suspended in Dulbecco's modified Eagle medium (DMEM, Thermo Fisher Scientific Inc.) containing 10 % fetal bovine serum (Thermo Fisher Scientific Inc.), were seeded at a density of 10,000 cells per 100 μL per well on a 96-well plate and incubated at 37 °C under 5 % CO_2_ for 2 days. After removing the medium, the cells were washed with PBS. Then, 100 μL of DMEM medium containing Aβ_40_ or Aβ_42_ aggregate (20 μM) was reaggregated for 12 h, and different concentrations of ADLumin-5 (0.1% DMSO) were incubated at 37 °C under 5 % CO2 for 24 h, then the cell viability was determined following the ATP assay method (CellTiter® 2.0 cell Viability Assay, Promega).

SH-SY5Y cells were also treated as above procedure, and the cell viability was determined following the MTT assay method.

**3D Spheroid Culture**

SH-SY5Y cells suspended in Dulbecco's modified Eagle medium (DMEM, Thermo Fisher Scientific Inc.) containing 10 % fetal bovine serum (Thermo Fisher Scientific Inc.), were seeded at a density of 10,000 cells per 200 μL per well on 96-well spheroid bottom plate and incubated at 37 °C under 5 % CO_2_ for 10 days, changing the medium every two days. After removing the medium, 100 μL of different concentrations of ADLumin-5 (0.1% DMSO) in the DMEM medium or DMEM medium containing Aβ_40_ or Aβ_42_ aggregate (20 μM) preaggregated for 12 h, and different concentrations of ADLumin-5 (0.1% DMSO) were added to each well, the plate was incubated at 37 °C under 5 % CO_2_ for 48 h. Cell viability was determined following the MTT assay method.

**Cell Uptake Study**

HMC3 cells were seeded in confocal culture dishes at a density of 1 × 10^4^ cells per well and incubated overnight in DMEM supplemented with 10% fetal bovine serum (FBS) at 37 °C in a humidified incubator with 5% CO_2_. Aβ^pH^ was prepared according to a previously reported method ^[3]^. Cells were divided into four treatment groups: Group 1: Aβ^pH^ (5 μM); Group 2: Aβ^pH^ (5 μM) + ADLumin-5 (10 μM); Group 3: Aβ^pH^ (5 μM) + ADLumin-5 (20 μM); Group 4: Aβ^pH^ (5 μM) + ADLumin-5 (20 μM) + BafA1 (100 nM). The indicated compounds were added to the corresponding confocal culture dishes and incubated for an additional 24 h at 37 °C under 5% CO_2_. After incubation, cells were washed with PBS. LysoTracker Red (100 nM) was then added to label lysosomes, and the cells were incubated at 37 °C for 2 h. Finally, cells were imaged using laser scanning confocal microscopy.

**Apoptosis Study**

SH-SY5Y cells were seeded at a density of 1 × 10^4^ cells per well in low attachment U-bottom 96-well plates to generate 3D spheroids. The culture medium was replaced every 2 days for 2 weeks to allow spheroid formation. Then, the 3D spheroids were treated with the following conditions: Aβ_40_ aggregates (20 μM); Aβ_42_ aggregates (20 μM); Aβ_40_ aggregates (20 μM) + ADLumin-5 (2 μM); Aβ_42_ aggregates (20 μM) + ADLumin-5 (2 μM). Fresh medium containing the indicated compounds was replaced every 2 days, for a total of two treatments. After the treatment, spheroids were collected and gently dissociated by pipetting to obtain a single-cell suspension. Apoptotic cells were then stained using the In Situ Cell Death Detection Kit, Fluorescein (Roche, Cat# 11684795910) according to the manufacturer’s instructions. Finally, apoptosis was quantified by flow cytometry using a SORP 5-laser BD LSRFortessa system.

Commercially available 100-day-old brain organoids were transferred to low-attachment 6-well plates and maintained in organoid culture medium for an additional 48 h before treatment. The organoids were then assigned to the following groups: Group 1: Aβ42 aggregates (5 μM); Group 2: Aβ42 aggregates (5 μM) + ADLumin-5 (2 μM); Group 3: ADLumin-5 (2 μM). Fresh medium containing the indicated compounds was replaced every 2 days, for a total of two treatments. Subsequently, organoids were collected and fixed with 4% paraformaldehyde (PFA). Single-cell suspensions were subsequently prepared from the organoids. Cells from each group were stained using the APO-DIRECT™ Flow Cytometry Kit according to the manufacturer’s instructions. Apoptosis was then quantified by flow cytometry using a SORP 5-laser BD LSRFortessa system.

**Assessment of Protein Degradation in Brain Homogenates**

To evaluate the in vivo Aβ degradation capacity of ADLumin-5, brain homogenates from wild-type (WT) mice were subjected to the following treatments: 1) Aβ_40_ aggregates (negative control); 2) Aβ_40_ aggregates + ADLumin-5 (positive control); 3) WT brain homogenate + ADLumin-5 (WT group); and 4) WT brain homogenate + Aβ_40_ aggregates + ADLumin-5 (in vivo simulation group). Before incubation with the primary antibody, signals from both the chemiluminescence (CL) and fluorescence (FL) channels were acquired using an IVIS imaging system. Subsequently, the samples underwent incubation with primary and secondary antibodies; following the secondary antibody incubation, CL and FL signals were acquired once again to facilitate comparative analysis. Based on estimations, the concentration of Aβ within this simulated sample falls within the physiologically relevant range of Aβ concentrations previously reported in the brain tissue of APP/PS1 transgenic mice (approximately 10–500 ng Aβ/mg brain tissue).

**Aβ Content in Three-Dimensional Brain Organoids**

Commercially available 100-day-old brain organoids were transferred to low-attachment 6-well plates and cultured for an additional 48 h. Organoids were then treated with Aβ_40_ aggregates (5 μM) alone or Aβ_40_ aggregates (5 μM) + ADLumin-5 (10 μM) for 2 days in organoid culture medium.

After treatment, organoids were collected and lysed on ice for 30 min using RIPA buffer. The lysates were diluted 1:10 and analyzed by ELISA. Aβ_40_ aggregate standards (6.25-800 ng/mL) were prepared in parallel to generate a standard curve.

Samples and standards were added to 96-well plates (100 μL per well) and incubated for 2 h at room temperature, followed by PBST washing and blocking with 2% BSA (1 h). Plates were then incubated with primary antibody (6E10, 100 nM) overnight at 4 °C, followed by HRP-conjugated goat anti-mouse secondary antibody (1:10,000, 1 h at room temperature). Signal was developed using TMB substrate and stopped with 0.18 M H_2_SO_4_, and absorbance was measured at 450 nm. Aβ40 concentrations in organoid samples were calculated from the standard curve.

**Ex Vivo Tissue Imaging**

Brains from 5×FAD mice were perfused, collected, and fixed overnight in 4% paraformaldehyde (PFA). After dehydration in 30% sucrose, 40 μm coronal sections were prepared using a cryostat and stored in PBS. Sections were washed with PBS, permeabilized with 0.1% Triton X-100 (10 min), and incubated in PBS containing 0.1% Triton X-100 and 2% BSA with ADLumin-5 (5 μM) for 48 h at room temperature in the dark with gentle shaking. Control sections were incubated under identical conditions without ADLumin-5. After washing with PBS, sections were stained with CRANAD-3 (1 μM) overnight at 4 °C, washed again, mounted with antifade mounting medium, and imaged using laser scanning confocal microscopy (CRANAD-3: Ex = 630-650 nm, Em = 660-740 nm).

To further investigate the structural and fluorescence properties of Aβ plaques, expansion microscopy (ExM) was performed on brain sections from 5×FAD mice. Briefly, 40 μm coronal sections were incubated overnight at 4 °C with Acryloyl-X, SE (AcX) to anchor proteins to the hydrogel matrix. Sections were then infiltrated with protein-retention ExM monomer solution containing acrylamide, sodium acrylate, crosslinker, and initiator, followed by in situ polymerization to form a tissue–hydrogel composite. After gelation, samples were digested with Proteinase K (1:100) at 37 °C for 6 h and expanded in ultrapure water to achieve an approximately 4× linear expansion.

Expanded gels were incubated with CRANAD-3 and ADLumin-5 (5 μM) overnight to allow probe penetration and binding to amyloid structures. Excess probes were removed by extensive washing in ultrapure water while maintaining the expanded state. Samples were then imaged using laser scanning confocal microscopy.

**In Vivo Imaging and Therapeutic Studies**

18-week-old female 5×FAD mice were randomly assigned to treatment groups (n = 5 per group), with wild-type mice (n = 5) used as controls. For in vivo chemiluminescence imaging, mice received intraperitoneal injections of ADLumin-5 (4 mg/kg; 1 mg/mL in 5% DMSO, 15% Cremophor, and 80% saline) on days 0, 68, 99, and 129. Chemiluminescence images were acquired every 5 min after injection using an IVIS imaging system, and semi-quantitative analysis was performed by measuring the average luminescence intensity within the brain region of interest (ROI).

For the treatment study, mice received intraperitoneal injections three times per week. The treatment groups were as follows:

Group 1 (wild-type control): ADLumin-5 (4 mg/kg, 80 μL; 1 mg/mL in 5% DMSO, 15% Cremophor, 80% saline).

Group 2 (5×FAD model control): Vehicle (80 μL; 5% DMSO, 15% Cremophor, 80% saline).

Group 3 (5×FAD treatment): ADLumin-5 (4 mg/kg, 80 μL; 1 mg/mL in 5% DMSO, 15% Cremophor, 80% saline).

**Ethics Statement**: All experimental protocols have been reviewed and approved by the Institutional Animal Care and Use Committee (IACUC) at Massachusetts General Hospital. (Approval No.: 2011N000161).

**ELISA Analysis of Aβ in Brain Tissue**

After treatment, mice were euthanized, and brain tissues were collected and lysed on ice for 30 min using RIPA lysis buffer. The lysates were diluted 1:10 and analyzed by ELISA. Aβ40 and Aβ42 aggregate standards (6.25-800 ng/mL) were prepared in parallel to generate standard curves.

Samples and standards (100 μL per well) were added to 96-well plates and incubated at room temperature for 2 h. After washing with PBST, the plates were blocked with 2% BSA for 1 h. Plates were then incubated overnight at 4 °C with primary antibody (6E10, 100 nM), followed by HRP-conjugated goat anti-mouse secondary antibody (1:10,000, 1 h at room temperature).

Signal was developed using TMB substrate, and the reaction was stopped with 0.18 M H_2_SO_4_. Absorbance was measured at 450 nm, and Aβ40 and Aβ42 concentrations in brain samples were calculated from the standard curves.

**Brain Tissue Staining**

Brains from the treatment mice were perfused, collected, and fixed overnight in 4% paraformaldehyde (PFA). After dehydration in 30% sucrose, 40 μm coronal sections were prepared using a cryostat and stored in PBS. Sections were washed with PBS and then stained with CRANAD-3 (1 μM) for 15 min at r.t, washed again, mounted with antifade mounting medium, and imaged using laser scanning confocal microscopy (CRANAD-3: Ex = 630-650 nm, Em = 660-740 nm)

**Experimental section**

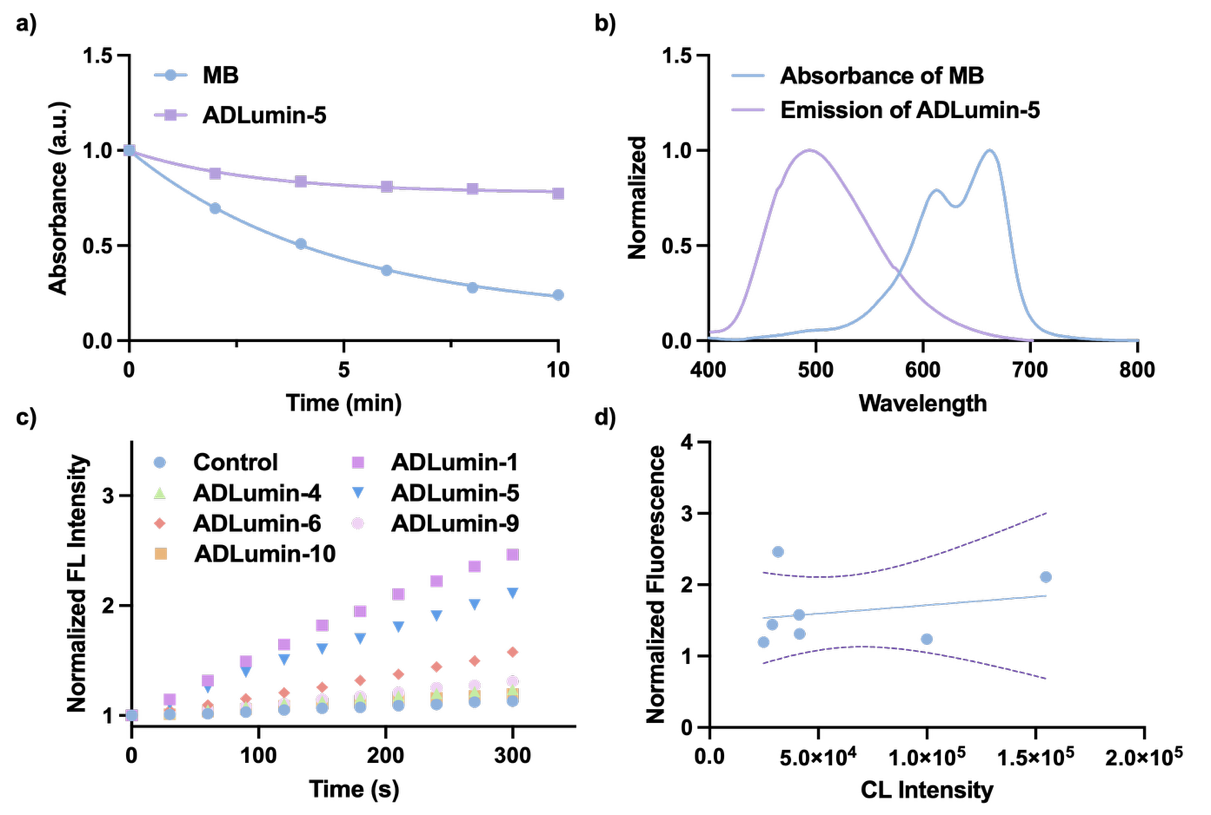

**Fig. S1 a)** The absorbance of MB or ADLumin-5 (25 μM) treated with ADPA (25 μM) after LED irradiation for different time points; **b)** Spectrum overlap of MB (absorbance) and ADLumin-5 (emission); **c)** Superoxide anion generation of ADLumin-Xs (25 μM) after being treated with DHE (25 μM); **d)** Correlation fitting of superoxide anion generation capability and chemiluminescence intensity for ADLumin-Xs.

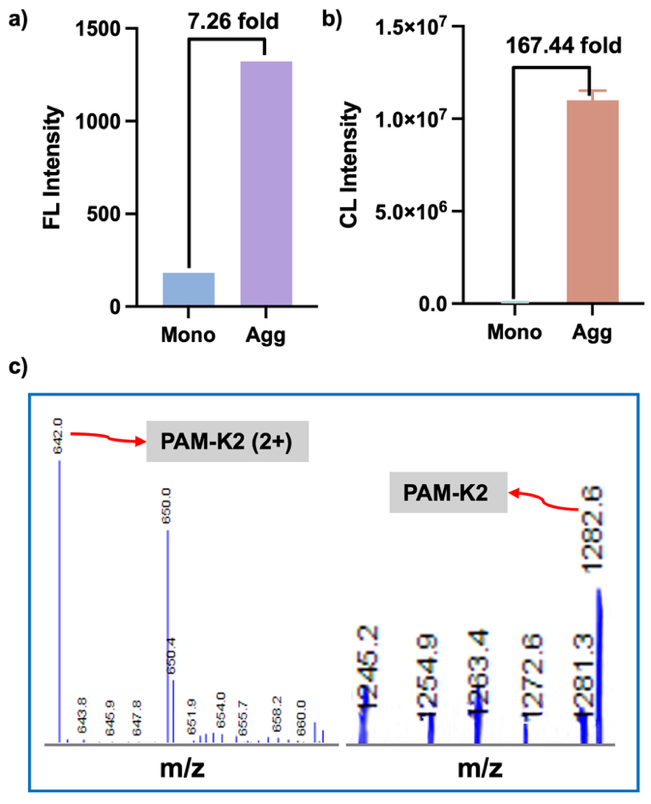

**Fig S2. a**) Fluorescence intensity quantification of PAM-K2 monomer or aggregate (5 μM) after adding Thioflavin T (0.5 μM) in PBS; **b**) Chemiluminescence intensity quantification of PAM-K2 monomer or aggregate (50 μM) with ADLumin-5 (50 μM); **c**) Mass spectrum of PAM-K2 with charge after photo-oxidation.

**Tab. S1 Oxidation efficiency of ADLumin-Xs on PAM-K2 aggregate**

| Compound | Ratio of oxygenated/non-oxygenated |
| --- | --- |
| ADLumin-1 | 50.0% |
| ADLumin-4 | 64.0% |
| ADLumin-5 | 77.1% |
| ADLumin-6 | 40.0% |
| ADLumin-9 | 20.0% |
| ADLumin-10 | 16.7% |
| ADLumin-15 | 30.0% |

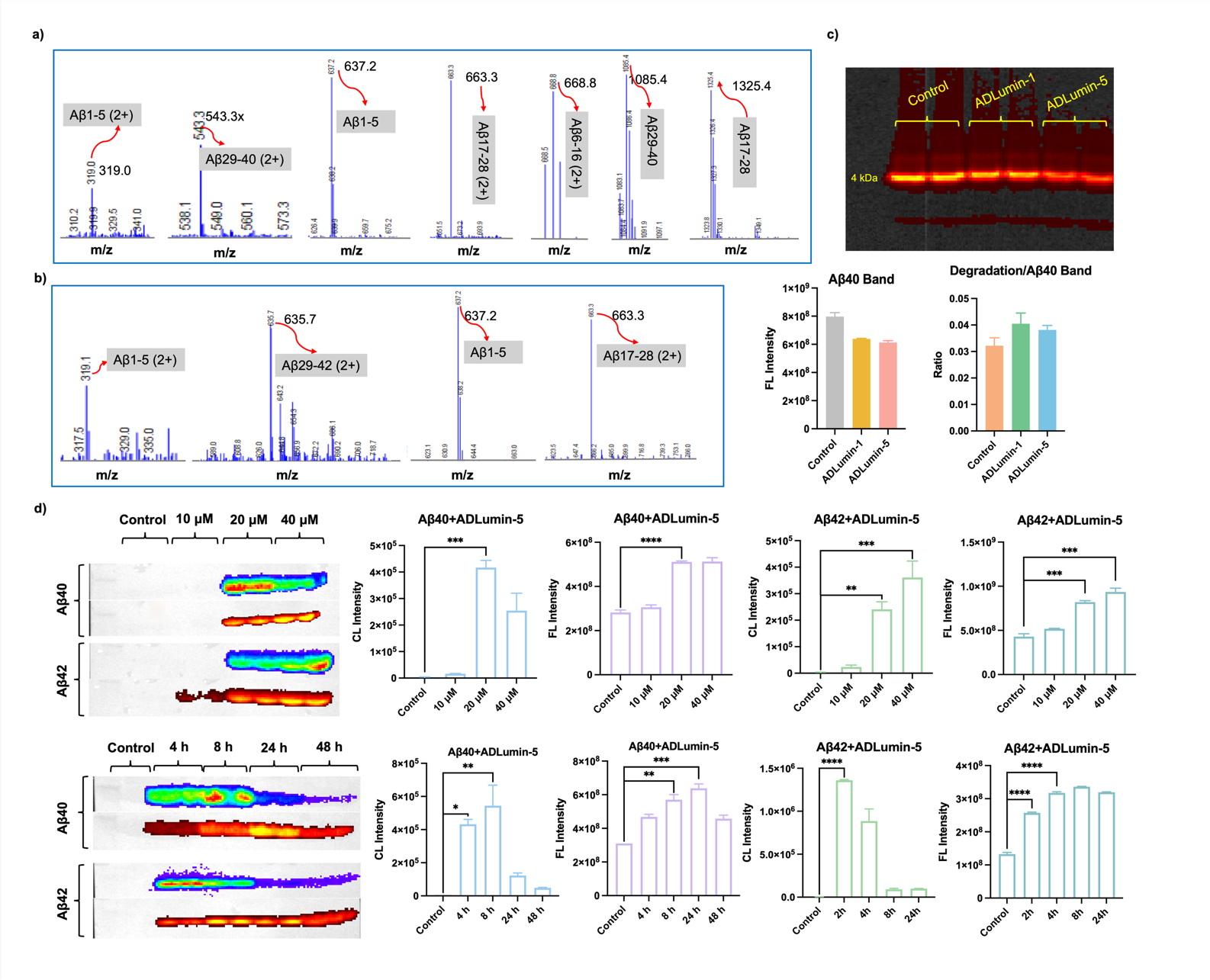

**Fig. S3** **a**) LC-MS analysis of Aβ_40_ incubated with ADLumin-5 after trypsin cleavage; **b**) LC-MS analysis of Aβ_42_ incubated with ADLumin-5 after trypsin cleavage; **c**) SDS-PAGE of FAM-Aβ_40_ aggregate incubated with ADLumin-Xs and quantification of monomer band and ratio of degradation to monomer; **d**) Western blot assay of Aβ ADLumin-5 for different concentrations or incubation times (before antibody incubation) and quantification of conjugation+degradation band in both chemiluminescence and fluorescence channels. Data are mean ± SD, n = 3. *p < 0.05, **p < 0.01, ***p < 0.001, ****p < 0.0001

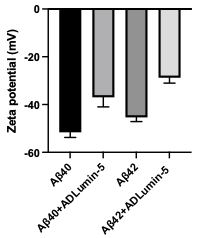

Fig. S4 Zeta potential changes of Aβ oligomers with or without ADLumin-5

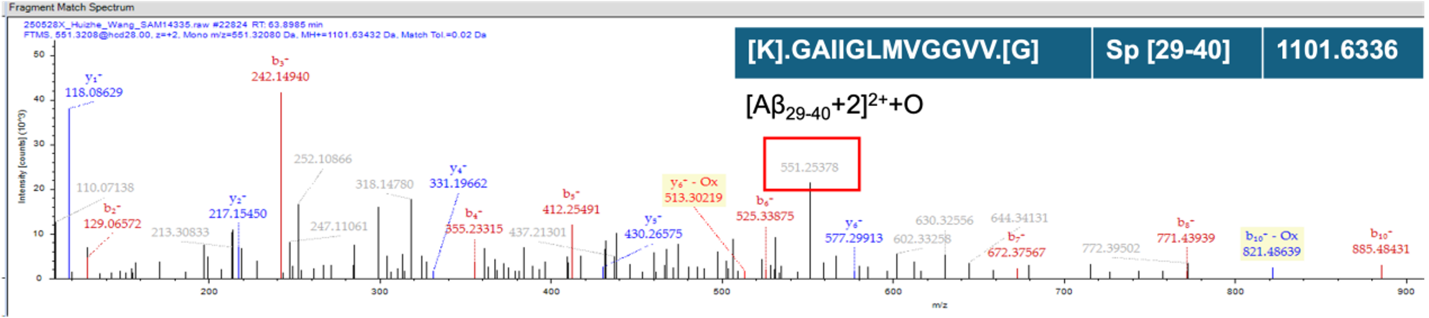

Fig. S5 Proteomics LC-MS analysis after incubation with ADLumin-5 and Aβ_40_

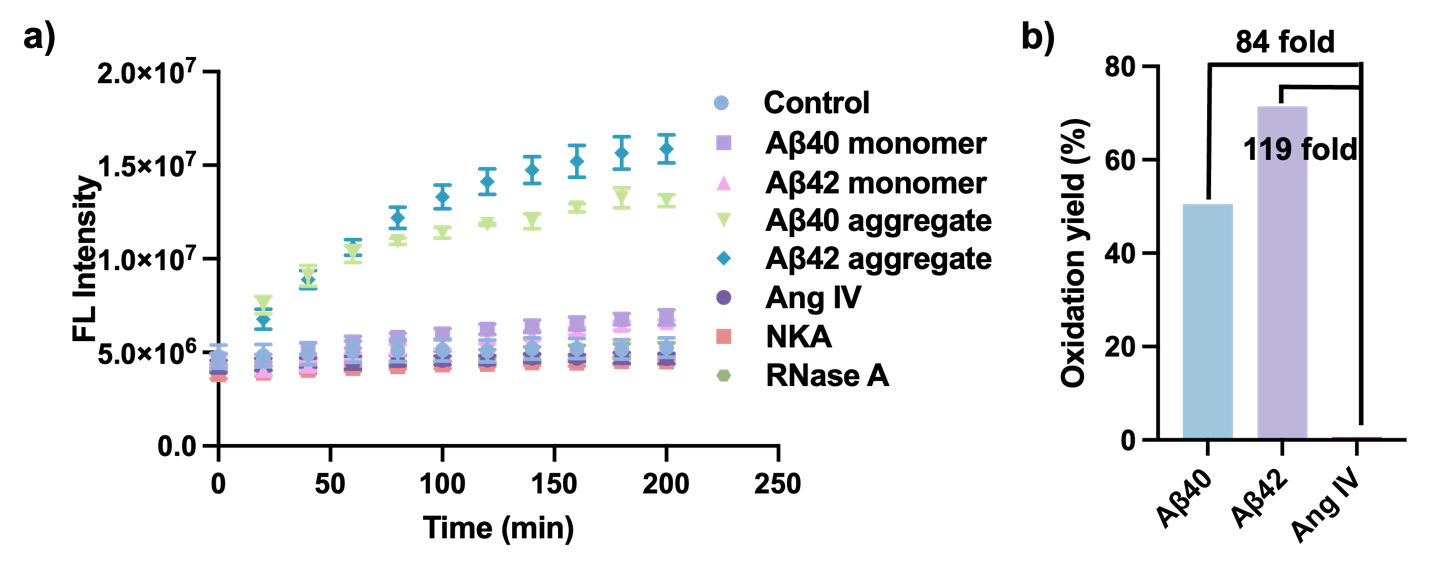

**Fig. S6 a)** Fluorescence intensity of Aβ_40_ or Aβ_42_ monomer and aggregate, and other non-β-sheet peptides with ADLumin-5 (2 μM); **b**) Oxidation efficiency of ADLumin-5 on Aβ and non-amyloid off-target model peptides containing oxidation-sensitive amino acid residues.

**Tab. S2 MS fragmentation ion information of the Aβ_40_ fragment (Aβ_29-40_)**

| #1 | b+ | Seq. | y+ | #2 |
| --- | --- | --- | --- | --- |
| 1 | 58.0672 | G | 1086.38 | 12 |
| 2 | 129.146 | A | 1029.32 | 11 |
| 3 | 242.306 | I | 958.246 | 10 |
| 4 | 355.465 | I | 845.087 | 9 |
| 5 | 412.517 | G | 731.927 | 8 |
| 6 | 525.677 | L | 674.875 | 7 |
| 7 | 656.875 | M | 561.716 | 6 |
| 8 | 756.008 | V | 430.517 | 5 |
| 9 | 813.06 | G | 331.384 | 4 |
| 10 | 870.112 | G | 274.332 | 3 |
| 11 | 969.244 | V | 217.28 | 2 |
| 12 | 1068.38 | V | 118.148 | 1 |

**Tab. S3 MS/MS fragmentation ion information of the Aβ_40_ fragment (Aβ_29-40_+O)**

| #1 | b+ | Seq. | y+ | #2 |
| --- | --- | --- | --- | --- |
| 1 | 58.0287 | G | 1102.38 | 12 |
| 2 | 129.066 | A | 1044.61 | 11 |
| 3 | 242.15 | I | 973.575 | 10 |
| 4 | 355.234 | I | 860.491 | 9 |
| 5 | 412.255 | G | 747.407 | 8 |
| 6 | 525.34 | L | 690.385 | 7 |
| 7 | 672.375 | M-Oxidation | 577.301 | 6 |
| 8 | 771.443 | V | 430.517 | 5 |
| 9 | 828.465 | G | 331.384 | 4 |
| 10 | 885.486 | G | 274.332 | 3 |
| 11 | 984.555 | V | 217.28 | 2 |
| 12 | 1084.38 | V | 118.148 | 1 |

**Tab. S4 Pearson colocation coefficient calculation**

|  | Pearson Correlation (r) | Mander M1 | Mander M2 |
| --- | --- | --- | --- |
| Control | 0.03 | 0.084 | 0.002 |
| ADLumin-5 (2 eq) | 0.30 | 0.424 | 0.306 |
| ADLumin-5 (4 eq) | 0.36 | 0.402 | 0.413 |
| ADLumin-5+BafA1 | 0.15 | 0.162 | 0.115 |

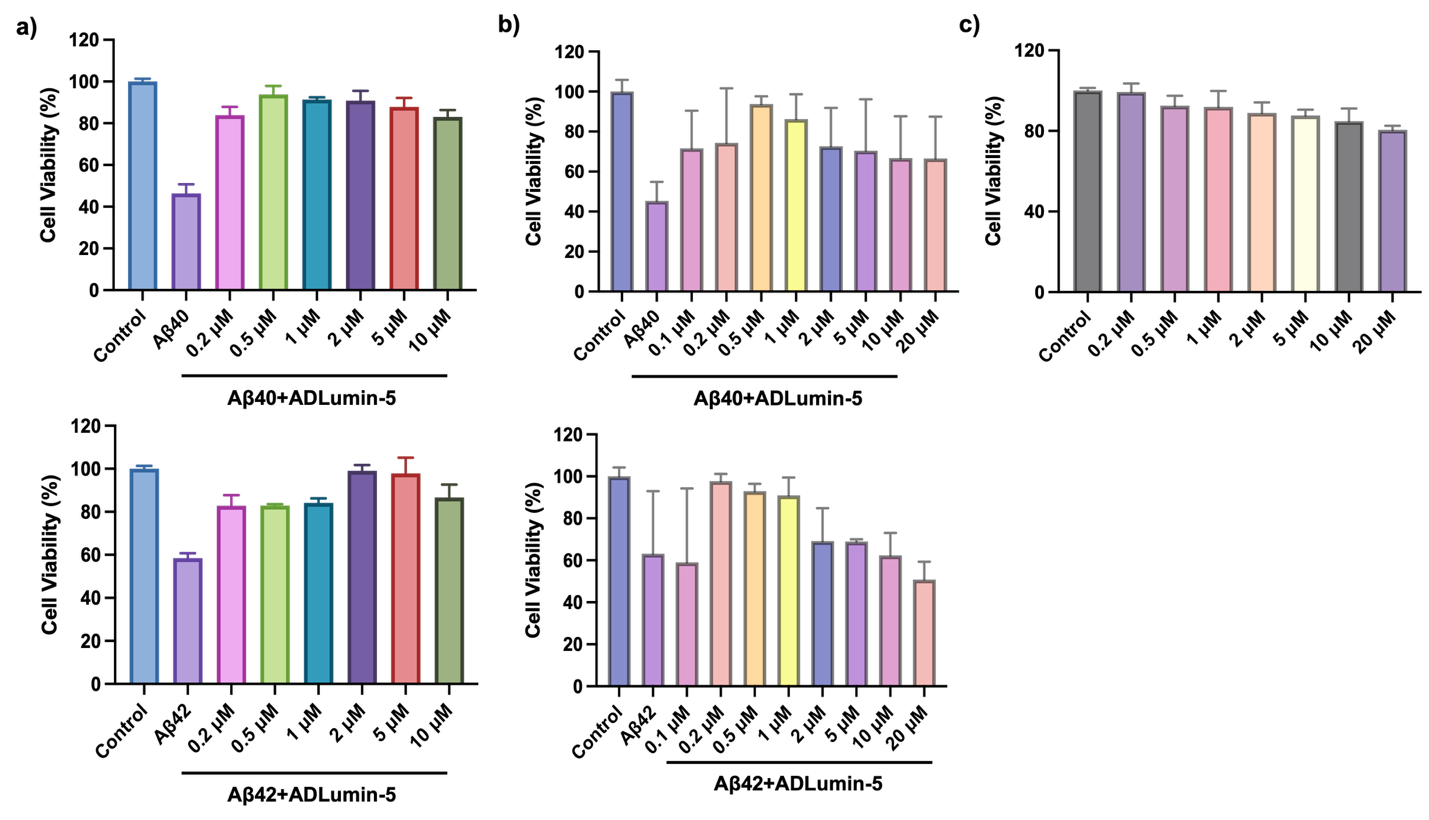

**Fig. S7** **a**) Cell viability assay by ATP kit of SH-SY5Y cells with different treatment Aβ (20 μM); **b**) Cell viability assessment with MTT method for 3D spheroids from SH-SY5Y cells; **c**) MTT assay of SH-SY5Y cells treated with different concentrations of ADLumin-5.

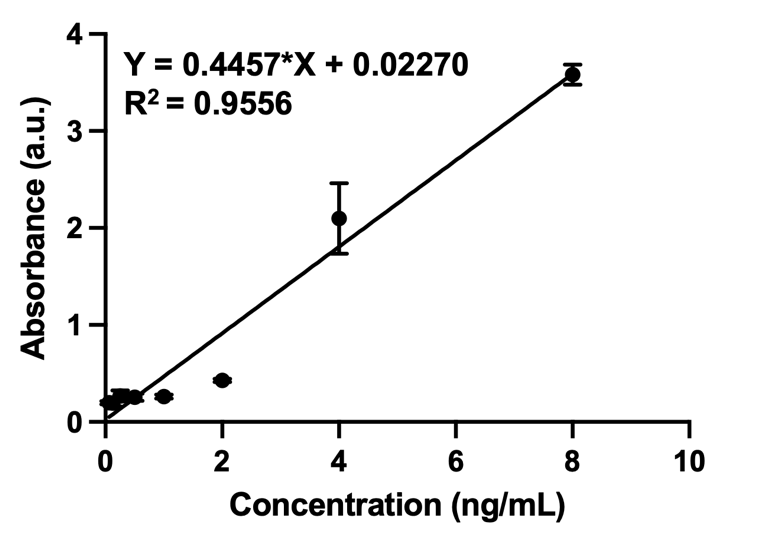

**Fig. S8** Aβ_40_ Standard Curve and Fitting Relationship

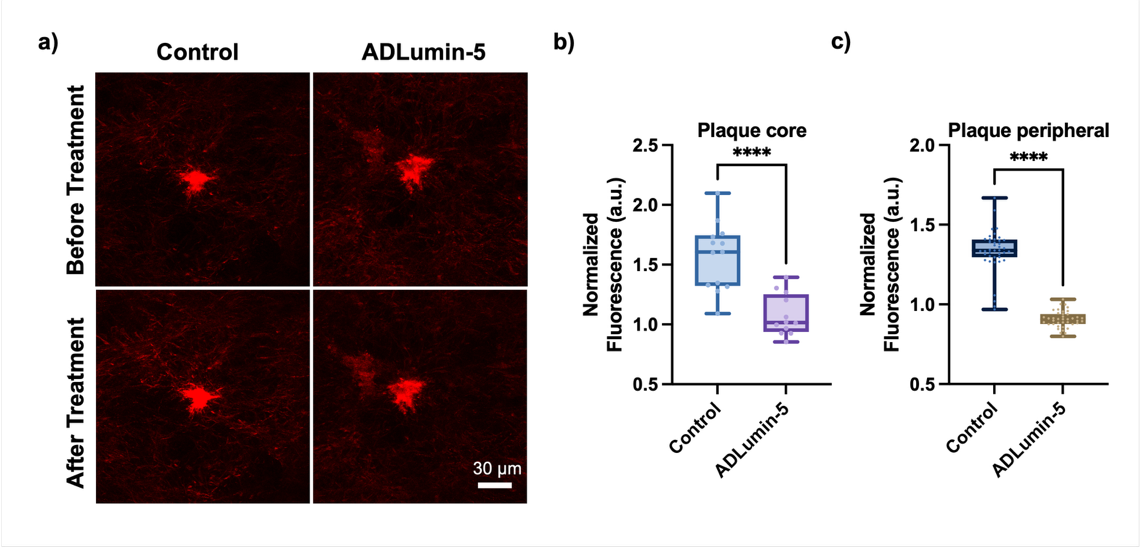

**Fig. S9 a**) Study of Aβ degradation in fixed mouse brain tissue treated or untreated with ADLumin-5 (not expanded sample); **b**) Normalized fluorescence intensity of plaque cores after treatment; **c**) Normalized fluorescence intensity of plaque periphery after treatment. Data are mean ± SD, n = 3. ****p < 0.0001

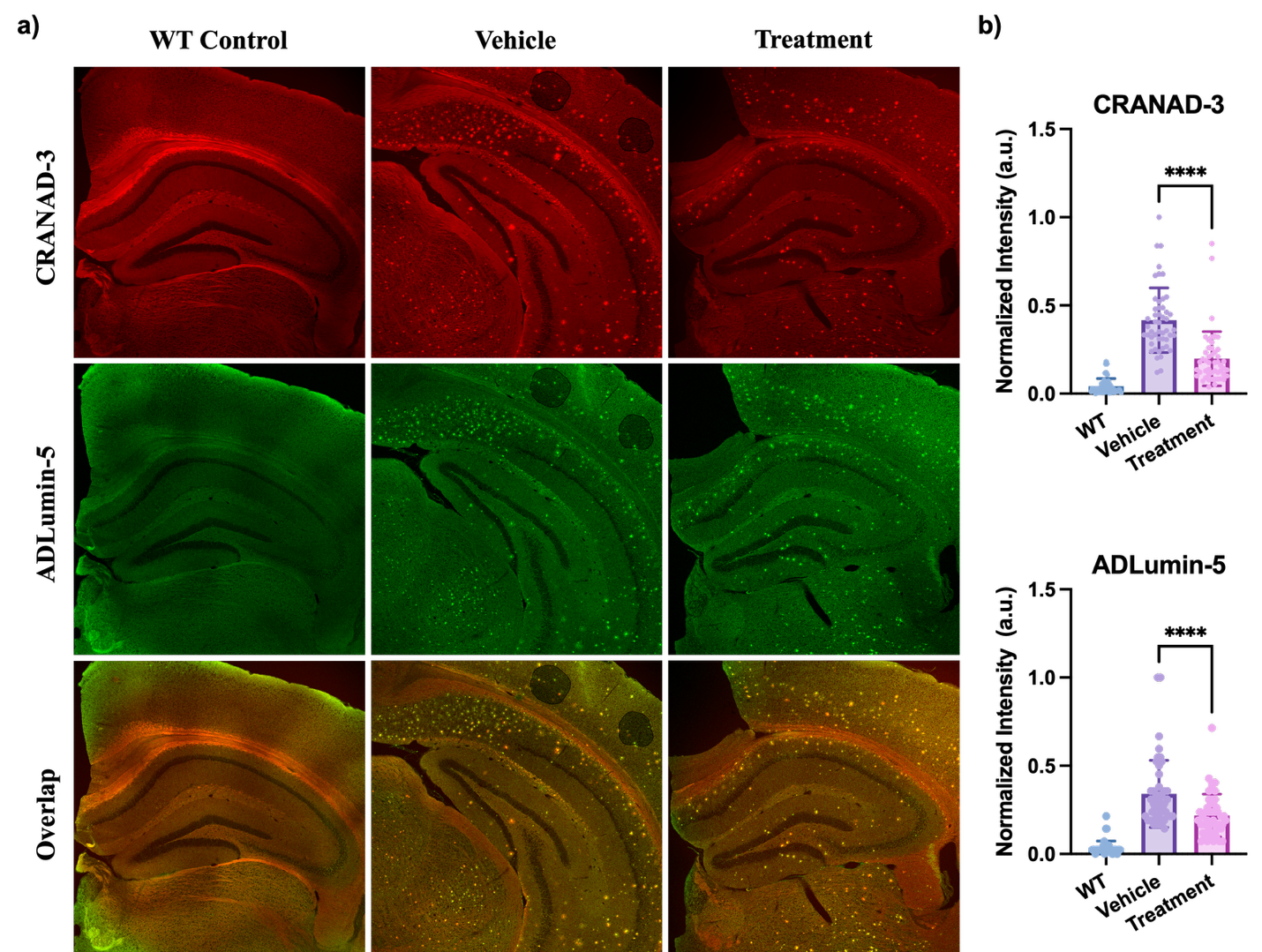

**Fig. S10**. Ex vivo fluorescence imaging and quantitative analysis of Aβ burden in mouse brain sections following ADLumin-5 treatment. Data are mean ± SD, n = 3. ***p < 0.001, ****p < 0.0001
